# Adenosine A2A and A2B receptor signaling in neurons promotes glucose and fatty acid release in the postprandial state

**DOI:** 10.64898/2026.05.29.728813

**Authors:** Daniel S. Lank, Brandon D. Seltzer, Stefan R. Hargett, Rahul M. Reddy, Julia Totten, Alexis J. Neimi, Michael C. Lemke, Mitchell E. Granade, Guzel K. Naik, Mark P. Beenhakker, Stephen B. G. Abbott, Joel Linden, Thurl E. Harris

## Abstract

Adenosine is a widely distributed signaling molecule whose levels rise during conditions of metabolic stress, hypoxia, or inflammation. Adenosine is a homeostatic regulator of neuronal, cardiovascular, immune, and metabolic functions through activation of adenosine receptors. Here, we show that administration of adenosine rapidly elicits an immediate and pronounced excursion of glucose and non-esterified fatty acids (NEFA) in mice refed for four hours but is greatly attenuated in fasted mice. This adenosine-mediated postprandial response suggests that adenosine is a potent regulator of postprandial nutrient handling. Selective agonists and antagonists of A2A and A2B adenosine receptors demonstrate that activation of either is sufficient to evoke adenosine’s metabolic response, but both receptors must be inhibited simultaneously to abolish it. Adenosine strongly stimulates hepatic glucose production and adipose lipolysis, and this catabolic activity depends on sympathetic nerve activity as it requires autonomic signal transmission and is inhibited by blocking adrenergic receptors. Genetic ablation studies identify A2A and A2B receptors expressed on neurons, likely central neurons, as the primary site of action for mediating adenosine’s effects on whole-body metabolism. Collectively, these data demonstrate that acute adenosine administration promotes centrally mediated metabolic effects, particularly during the postprandial period. Rigorous dissection of signaling pathways shows that A2A and A2B receptors are individually sufficient and collectively necessary for adenosine-mediated glucose and fatty acid excursion.

## INTRODUCTION

The postprandial state refers to a period several hours after a meal during which nutrients are digested, absorbed, and stored, typically encompassing early glucose excursions over ∼2–4 hours and more prolonged lipid (NEFA/triglyceride) excursions over ∼6–8 hours after fat-rich meals [1–3]. Postprandial increases in circulating glucose and fatty acids are normally resolved as insulin promotes glycogen storage in liver and triglyceride storage in adipose tissue [3]. In modern Western society adults spend most of the waking day in a postprandial or “fed” state, making this the dominant metabolic condition in which fuel handling is regulated and a potentially critical point for dysregulation [2]. The transition from fasting to postprandial states involves a rapid shift from catabolic signaling (glucagon, catecholamines, cortisol) to anabolic insulin signaling. Disruption to this balance contributes to metabolic disease, as seen in obesity-associated insulin resistance with chronically elevated glucose and lipids [1].

Adenosine is a purinergic signaling molecule that has a plethora of different receptor-mediated effects across the body. Its production is generally associated with injury, inflammation, metabolic stress, hypoxia, ischemia, and other pathophysiological conditions [4]. Extracellular adenosine arises primarily through enzymatic breakdown of cellular ATP. Under conditions of cellular stress or increased metabolic demand, intracellular ATP is hydrolyzed faster than it can be replaced, causing intracellular adenosine levels to increase. This accumulated adenosine can then pass down its concentration gradient into the extracellular space via equilibrative nucleoside transporters (ENTs). Adenosine can also be derived from ATP leakage into the extracellular space where ectonucleotidases CD39 and CD73 rapidly metabolize ATP into adenosine [4,5]. Collectively, these processes lead to elevated extracellular adenosine and subsequent adenosine signaling. Once produced, adenosine is rapidly metabolized by adenosine deaminase into inosine, or can be phosphorylated into 5’ AMP by adenosine kinase [6]. Due to the abundance of these enzymes, adenosine has a very short half-life (1-2 seconds) [7,8] and is restricted to autocrine and paracrine signaling.

Adenosine signals via a family of GPCRs known as adenosine receptors. There are four in total: A1 and A3 which couple to the G_αI/o_, A2A which couples to the G_αs_, and A2B which couples to both G_αs_ and G_αq_ subunits [9]. In the periphery, adenosine is involved in a wide variety of functions ranging from modulating inflammation, lowering heart rate, and stimulating vasodilation to name a few. In the central nervous system (CNS) both A1 and A2A receptors are necessary for modulating the sleep/wake cycle and can modulate autonomic signaling [4,6,9].

Adenosine’s contribution to metabolic regulation remains incompletely understood. It is well-established that acute adenosine signaling via the A1 receptor potently suppresses adipocyte lipolysis and reduces circulating non-esterified fatty acids (NEFA) [10–12], and the A3 receptor modestly regulates liver glucose output [13]. The roles of A2A and A2B on metabolism, however, are less clear. Some groups have suggested that activation of these two receptors lowers glucose and increases insulin sensitivity, while others report that they promote insulin resistance and increase circulating glucose levels. For example, Pfeiffer and colleagues have shown that chronic agonism of A2A and A2B in obese mice increases energy expenditure, reduces body weight, and improves whole-body glucose tolerance and insulin sensitivity [14,15]. Verman et al. 2025 showed that fat-specific deletion of A2A worsened diet-induced obesity, indicating an anti-diabetic role for A2A [16]. Contrarily, Csóka et al. 2014 showed that deletion of the A2B receptor reduced adipose tissue inflammation and improved whole-body insulin sensitivity [17]. Figler et al. 2011 showed that acute administration of the non-hydrolysable adenosine analogue 5-(N-ethylcarboxamido) adenosine (NECA) to lean mice decreased glucose tolerance in an A2B-dependent manner, and that acute dosing of a A2B antagonist improved insulin sensitivity in obese mice [18]. Sacramento and Conda 2020 showed that chronic antagonism of A2A and A2B increased insulin sensitivity in rats fed a high sucrose diet, but paradoxically decreased insulin sensitivity in lean rats [19]. Most of these studies implement the use of small molecule agonists and antagonists to selectively activate or inhibit adenosine receptors in a targeted way, with few studies examining metabolism when adenosine itself is administered *in vivo*. One notable exception is Tadaishi et al. 2018, which showed that acute adenosine administration caused a dose dependent glucose excursion, but did not specify which receptor was responsible [20]. These studies show that the metabolic effects of adenosine receptor activation are highly context-dependent, but collectively support the conclusion that modulation of adenosine signaling is sufficient to affect whole-body metabolism and may serve as a favorable target for mitigating metabolic diseases like obesity and diabetes.

Here we propose a novel role for central adenosine signaling in modulating peripheral metabolism. We show that administration of a single high dose of adenosine in mice is sufficient to promote an acute and drastic increase in circulating blood glucose and non-esterified fatty acids (NEFA), particularly when it is administered in the postprandial state. Using small molecule adenosine receptor agonists and antagonists, we determined that both A2A and A2B receptors mediate these effects. Adenosine-mediated glucose and NEFA excursion can be blocked by inhibiting α and β adrenergic receptors. Finally, we found that genetic deletion of both A2A and A2B receptors in neurons was sufficient to block the adenosine effect on peripheral metabolism. We conclude that A2A and A2B signaling in the CNS can modulate peripheral metabolism via sympathetic signaling.

## METHODS

### Materials

All small molecules were purchased from Cayman Chemical (Ann Arbor, MI) unless otherwise specified. The A2A receptor agonist ATL-313, the A2B receptor antagonist ATL-844, and the pan adenosine antagonist BW-1433 were generously provided by Adovate, LLC. (Charlottesville, VA). ^3^H 2-deoxyglucose was purchased from PerkinElmer (Waltham, MA).

### Mice

All animals were bred and maintained in accordance with the University of Virginia Animal Care and Use Committee regulations and the study was approved by the ACUC ethics committee. Unless otherwise specified, 12-18 week old male mice were used for all experiments. Mice containing floxed alleles of *Adora2a* and *Adora2b* have been previously described [21–24], and were backcrossed for more than five generations onto the C57Bl6/J line. Cre-expressing mouse lines were obtained from JAX and crossed into the floxed Adora lines, specifically AdipoQ-Cre/ERT2 (JAX #02524), Baf-53b-Cre/Actl6b-Cre (JAX #027826), ROSA26-Cre/ERT2 (JAX#008463), and UBC-Cre/ERT2 (JAX #007001).

### Adeno-Associated Virus (AAV)

AAV serotype 8 vectors were obtained from Addgene (Watertown, MA) and carried the thyroxine-binding globulin (TBG) promoter driving either Cre (Addgene #107787-AAV8) or eGFP (Addgene #105535-AAV8). For liver expression, ten to twelve-week old male mice were tail vein injected with 2.5 x 10^10^ GC/mouse. All mice were given a three-week washout period before subsequent experiments were conducted.

The AAV for peripheral neuron expression of Cre was made at VectorBuilder (Chicago, IL). The AAV9-PHP.S virus was chosen based on its previously demonstrated penetrance into peripheral neurons, while avoiding crossing the blood brain barrier [25]. The expression vector was synthesized where the SYN1 promoter drives EGFP:T2A:Cre expression module (pAAV[Exp]-SYN1>EGFP(ns):T2A:Cre:WPRE), and then packaged into AAV using the PHP.S serotype. Control virus for peripheral neuron expression was pAAV-PHP.S-CAG-tdTomato (Addgene #59462-PHP.S). UBC-A2A-A2B ^flox/flox^ Cre – mice were given AAV9-PHP.S-hSyn1-Cre-eGFP or AAV9-PHPS-CAG-tdTomato when the mice were 14-16 weeks old. Both viruses were administered at a dose of 1×10^12^ GC/mouse. All mice were given a three-week washout period before subsequent experiments were conducted.

### Tamoxifen treatment

Tamoxifen was dissolved in 100% peanut oil (Sigma Aldrich, Cat. No. P2144) at a concentration of 20 mg/ml. Eight- to twelve-week-old mice were given daily i.p. injections of 100 µL of tamoxifen (2 mg/mouse/day) for five consecutive days. Mice were then given a two-week washout period before subsequent experiments were performed.

### Fasting and refeeding experiments

Animals were placed in fresh Sani-chip cages without food, but access to water, at 5 PM the day preceding the experiment. At 7 AM the following morning, mice were either given *ad libitum* access to chow food pellets (refed) or continued fasting for 4 additional hours (fasted). At 11 AM, the fasted and refed mice were subjected to experimentation.

### Quantification of blood glucose, NEFA, and hormones

Blood glucose was measured from the tail vein using a One-Touch Ultra glucometer using Unistrip1 test strips (UniStrip Technologies Cat. No. 24850). Blood was collected from the tail vein into serum separation tubes (Greiner, Cat. No. 450472), allowed to coagulate on ice for 30 minutes, then the centrifuged at 4,500 x g for 10 minutes to separate out blood serum. Samples were stored at -80°C until analysis. Serum NEFA was measured using the NEFA-HR (2) Wako Diagnostics Kit (Cat. No. 994-91801). Insulin was measured using the Crystal Chem Ultra-Sensitive Mouse Insulin ELISA Kit (Cat. No. 62100).

### Dissolution of adenosine

Powdered adenosine was purchased from ThermoFisher (Cat. No. A10781.09). Adenosine was freshly prepared for each experiment. Powdered adenosine was dissolved in saline with 5% Kolliphor EL (Sigma Aldrich, Cat. No C5135) to a final concentration of 10 mg/mL. This solution was then heated to 55°C for 10 minutes, vortexed vigorously, then incubated for an additional 10 minutes at 55°C. After a second bout of vigorous vortexing, suspended adenosine was ready for injection.

### Quantification of liver glycogen was done via Amplex Red assay

Liver glycogen was measured using the AmplexRed assay. Briefly, 100 mg of liver tissue was homogenized 1 mL H_2_O. Samples were heated at 70°C for 15 minutes, then centrifuged at 17,000 x G for 10 minutes. The supernatant was then incubated with (+AMG) or without (-AMG) amyloglucosidase (1 mg/mL) in 0.2 M NaOAC (pH 4.8) for 1 hour at 37°C. 20 µL of each sample lysate was then incubated in a 96 well plate with 100 µL of horseradish peroxidase (0.5 U/mL), glucose oxidase (0.25 U/mL), FAD (10 µM), EDTA (1 mM), MgCl_2_ (1 mM), KH_2_PO_4_ (100 mM, pH 6.8), and AmplexRed (20 mM) for 10 minutes in the dark at room temperature. Absorbance was measured at 587 nm after an initial excitation at 535 nm. Values of the -AMG treatment were subtracted from the +AMG to remove background free glucose. All values were normalized to total protein concentration.

### Genomic DNA extraction and PCR

Genomic DNA was extracted using the Invitrogen PureLink Genomic DNA Mini Kit (Cat. No. K182001). The isolated gDNA was then amplified in a PCR reaction implementing primers we designed to flank the excised portion of A2A and A2B genes. Primers used were A2A Forward (CTCTCCTTTGTCTGCCCTGA), A2A Reverse (GACAGAGACGAGGAGAGGC), A2B Forward (ATTAGGGGACTGCAGGTGTG), A2B Reverse (ACGAAGTGCA AGAAGCCAGT).

### Western blotting and quantitative analysis

Protein lysates from adipose and liver tissue were extracted via Potter-Elvehjem homogenization. Briefly, ∼100 mg of tissue was homogenized in lysis buffer (50 mM Tris Base pH 7.4, 10 mM NaF, 1 mM EDTA, and 1% Triton-X) supplemented with protease and phosphatase inhibitors. Samples were then centrifuged at 17,000 x g for 10 minutes at 4°C, and supernatant was taken. Protein concentration was quantified via Pierce BCA assay (Thermo Fisher, Cat. No. 23225). Protein samples were all diluted to an identical concentration in 1X Laemmli buffer and heated at 70°C for 20 minutes. 30 µg of protein sample was loaded to each lane, ran in an 10% acrylamide SDS-PAGE gel, and transferred onto a PVDF membrane. Membranes were blocked in 10% dry milk in TBST for 1 hour, incubated in primary antibody overnight, and then blotted with secondary antibody for 1 hour before imaging. Membranes were probed with vinculin (Cell Signaling, #13901) and phospho-CREB (Cell Signaling, #9198) antibodies.

### [^3^H] 2-DG Experiments

2-deoxy-D-[1,2-3H]-glucose ([^3^H] 2-DG) was purchased from PerkinElerm (Waltham, MA). Mice were fasted and refed as previously described. Five refed mice were injected with 10 µCi of ^3^H 2-DG and vehicle (5% kolliphor in saline) and five mice were injected 10 µCi of ^3^H 2-DG and adenosine (100 mg/kg). Over two hours, blood was collected at 0, 15, 30, 60, and 120 minutes, and serum was extracted as previously described. After 120 minutes, the animals were sacrificed and tissues were harvested and stored at -80° C until processing. Quantification of ^3^H 2-DG from serum was performed as previously described [26]. Briefly, 3 µL of serum was added to 200 µL of 3.5% perchloric acid and centrifuged at 14,000 x g for 10 minutes to pellet protein. 180 µL of the supernatant was combined with 50 µL of 2.2 M HKCO_3_ to neutralize the supernatant. From this mixture, 200 µL was added to 4 mL of scintillation fluid (Optiphase Hisafe 3) and measured in a scintillation counter (Beckman-Coulter, LS 6500). The decays per minute (dpm) were normalized to the corresponding µmol of cold glucose at each time interval to yield specific activity (dpm/ µmol glucose).

### Intracerebroventricular injections

Intracerebroventricular (i.c.v.) injections were performed using the Neurostar robotic stereotaxic system and a 34-gauge NanoFil needle (Cat. No. NF34BV). 15-18 week old WT male mice were anaesthetized with isoflurane before the beginning of the procedure. Following exposure of the skull, a small burr hole was drilled at the injection site, and injections were targeted to the lateral ventricle using the following coordinates relative to bregma: anterior–posterior (AP) −0.7 mm, medial–lateral (ML) −1.1 mm, and dorsal–ventral (DV) −2.0 mm. Vehicle (10 % kolliphor in saline) or BAY-60-6583 was infused at a rate of 0.2 µL/min using StereoDrive software. Following the infusion, the needle was left in place for 5 min to minimize reflux before slow withdrawal.

## RESULTS

### Adenosine administration promotes an increase in glucose and fatty acids

CCPA, an agonist selective for the A1 adenosine receptor, displays differential responses depending on the metabolic state [27]. To explore this further, we conducted experiments examining the effects of adenosine administration in mice under fasted and refed conditions. Wild-type C57Bl6/J (WT) mice were fasted overnight then subsequently given *ad libitum* access to food (refed) or continued fasting (fasted) for 4 hours the following morning. After this 4-hour window, all mice were given an intraperitoneal (i.p.) injection of adenosine (100 mg/kg), and their blood glucose and serum NEFA recorded over a two-hour period. Interestingly, while fasted mice displayed a moderate glucose excursion and essentially no NEFA excursion compared to vehicle-treated controls (Fig. 1 A, E), refed mice given adenosine displayed an immediate and profound induction in glucose and NEFA compared to vehicle-treated controls (Fig. 1 B, F). Direct comparison of refed and fasted adenosine-injected mice illustrates the significant differences in glucose (Fig. 1 C, D) and NEFA (Fig. 1 G, H) that accompanies these difference metabolic states.

**Figure 1:**
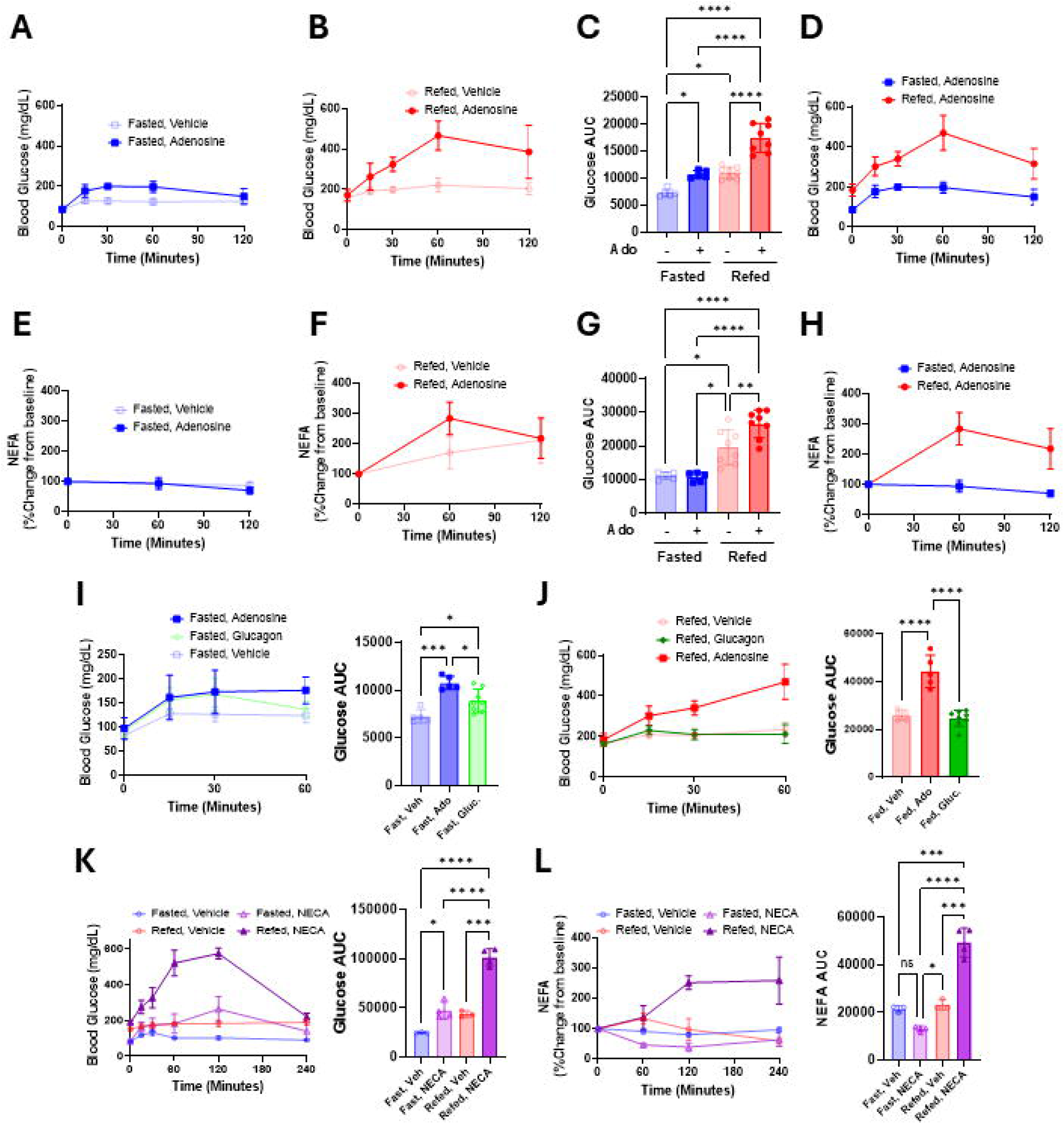
Adenosine administration causes a profound and acute increase in blood glucose and serum NEFA in the postprandial state: WT mice were fasted and/or refed and injected with an i.p. bolus of either vehicle or adenosine (100 mg/kg), and blood glucose and serum NEFA was measured over a two-hour period. Serum NEFA is reported as a percent change in reference to the t = 0 reading. **(A)** Blood glucose values from vehicle and adenosine-treated fasted and **(B)** refed mice. (**C**) Comparison of glucose excursion area under the curve (AUC) between 1A and 1B. (**D**) Comparison of glucose excursions between 1A and 1B. (**E)** Serum NEFA levels of vehicle and adenosine-treated fasted and **(F)** refed mice. (**G)** Comparison of serum NEFA excursion AUC between 1E and 1F. **(H)** Comparison of serum NEFA excursion from 1E and 1F. WT mice were injected i.p. with vehicle, glucagon (0.02 mg/kg) or adenosine (100 mg/kg) and blood glucose was measured over a one-hour period in **(I)** fasted and **(J)** refed mice. WT fasted and refed mice were injected i.p. with the pan-adenosine receptor agonist NECA (0.02 mg/kg) and their **(K)** blood glucose and **(L)** serum NEFA were measured over a four-hour period. N’s for each group range from 4-8. Error bars represent SD. *p < 0.05, **p < 0.01, ***p < 0.001, ****p < 0.0001. All statistical analyses were done by one-way ANOVA (I-J) or two-way ANOVA (A-H, K-L).

To gain an appreciation for the magnitude of glucose excursion stimulated by adenosine, we compared it with glucagon, an endogenous hormone with a well-characterized ability to increase blood glucose. When fasted mice were given a bolus of glucagon (20 µg/kg) the resulting glucose release was similar to adenosine (Fig. 1 I). However, when refed mice were given the same dose of glucagon, the glucose release was significantly lower than adenosine-mediated glucose release (Fig 1 J).

NECA is a pan adenosine receptor agonist with a higher affinity than adenosine, a much longer half-life, it cannot be metabolized into any of the adenosine metabolites [28]. Mice were fasted and refed as previously described and given an i.p. bolus of NECA at 20 µg/kg. Similar to adenosine, NECA was sufficient to produce a mild glucose induction in fasted mice and a pronounced glucose induction in refed mice (Fig. 1 K). NECA also promoted a profound increase in serum NEFA levels in refed mice, and a modest decline in fasted mice (Fig. 1 L).

To investigate if there were any sex-specific differences in adenosine’s ability to stimulate glucose and NEFA excursions, we subjected a cohort of female WT C57BL/6 mice to adenosine treatment under fasted and refed conditions. We found that fasted female mice did not produce a significantly different glucose or NEFA excursion comparted with vehicle-treated controls (Sup. Fig. 1 A, D). However, like male mice, refed female mice did display increased glucose and NEFA excursion in response to an adenosine challenge (Sup Fig. 1 B, E). Collectively, these data indicate that adenosine can acutely and profoundly promote glucose and fatty acid excursion, which is enhanced under postprandial conditions in both male and female mice. Moreover, adenosine elicits a glucose response similar to that of the counter regulatory hormone glucagon under fated conditions, but under postprandial conditions elicits a much larger glucose excursion.

### Adenosine-mediated glucose and NEFA release involves A2A and A2B signaling

To investigate which of the four adenosine receptors is responsible for mediating these metabolic effects, we examined the effect of acute administration of various selective agonists to individually stimulate each receptor under both fasted and refed conditions. Mice were given an i.p. bolus of either the A1-selective agonist CCPA (0.5 mg/kg), A2A-selective agonist ATL-313 (0.2 mg/kg), A2B-selective agonist BAY-60-6583 (0.2 mg/kg), or the A3-slective agonist 2-Cl-IB-MECA (0.2 mg/kg) under fasted and refed conditions. These agonists all have a substantially longer half-life than adenosine itself and have much higher affinity for their respective receptor than adenosine allowing for significantly lower doses to be used [29,30]. Agonism of A1 showed no significant effect on glucose excursion (Fig 2. A), however, we did observe suppressed lipolysis in the fasted, but not refed mice (Fig. 2 B). This is consistent with the well-established role of A1 signaling in adipose tissue to suppress lipolysis, and that A1 activity is potentiated in adipose tissue under fasted conditions [7–9, 22]. A2A agonism was sufficient to stimulate a glucose and NEFA excursion, which was significantly higher in refed animals (Fig. 2 C, D). A2B agonism was also sufficient to stimulate an increase in glucose and serum NEFA and was likewise potentiated in refed animals (Fig. 2 E, F). A3 agonism had no effect on glucose or NEFA in either fasted or refed state (Fig. 2 G, H). These data show that either of the G_αs_ coupled adenosine receptors, A2A and A2B, are sufficient to stimulate glucose and NEFA release, that is potentiated under postprandial conditions. The G_αi_ coupled adenosine receptors A3 and A1 do not appear to affect postprandial glucose, and only A1 signaling reduced NEFA under fasted conditions.

**Figure 2:**
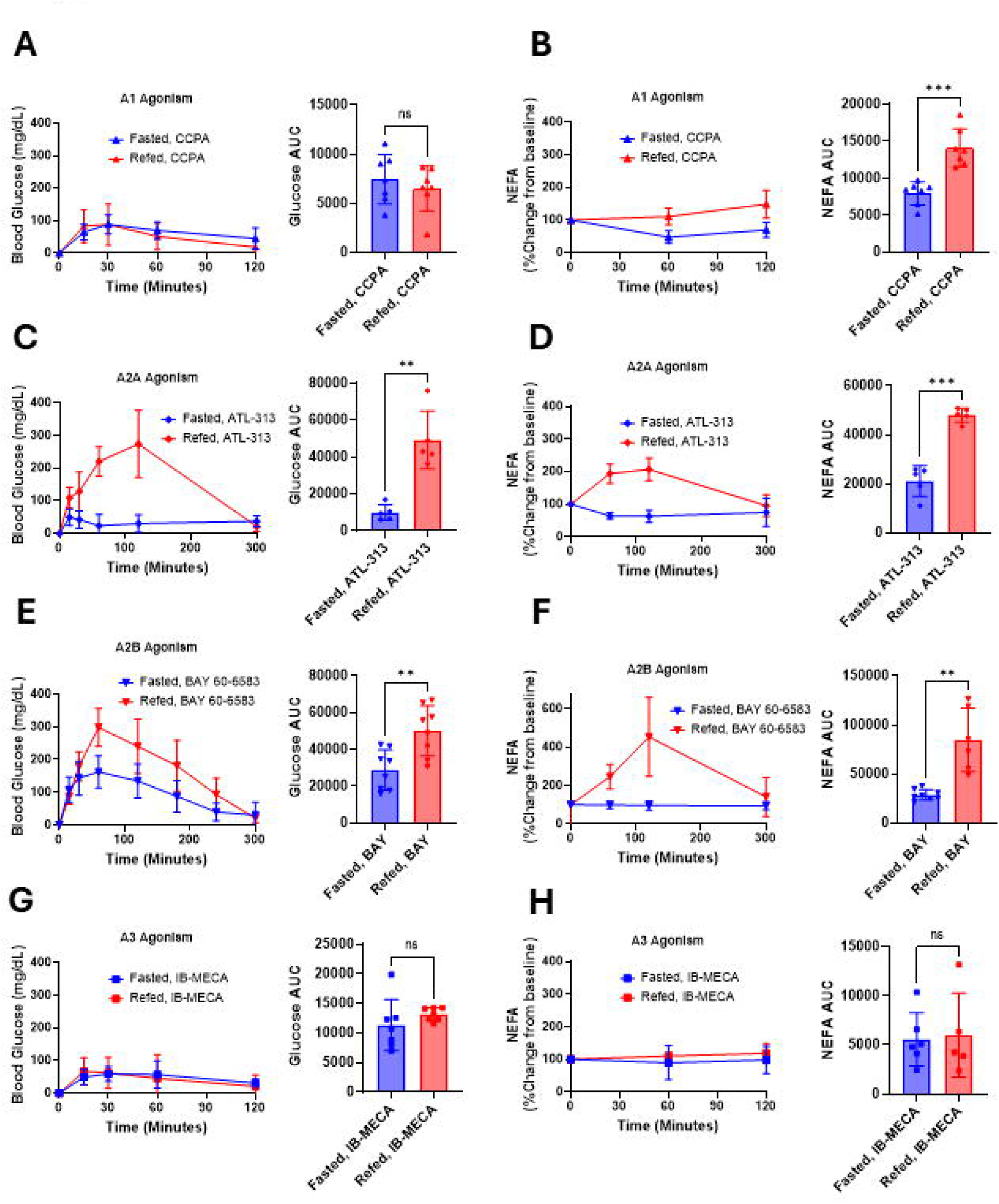
Postprandial agonism of either A2A or A2B receptors is sufficient to promote a glucose and serum NEFA excursion. Fasted and refed mice were challenged with an i.p. bolus of different small molecule agonists specific to each adenosine receptors (A1, A2A, A3, and A2B). Blood glucose and serum NEFA was measured throughout the experiment. To account for intrinsic differences in blood glucose and serum NEFA between fasted and refed groups, blood glucose is reported in reference to the t = 0 reading, and NEFA is reported as percent change from t = 0 levels. **(A)** Blood glucose and **(B)** serum NEFA changes in response to A1 agonist CCPA (0.5 mg/kg). **(C)** Blood glucose and **(D)** serum NEFA changes in response to A2A agonist ATL-313 (0.2 mg/kg). **(E)** Blood glucose and **(F)** serum NEFA changes in response to A2B agonist BAY-60-6583 (0.2 mg/kg). **(G)** Blood glucose and **(H)** serum NEFA changes in response to A3 agonist IB-MECA (0.2 mg/kg). N’s for each group range from 5-8. Error bars represent SD. *p < 0.05, **p < 0.01, ***p < 0.001, ****p < 0.0001. All statistical analyses were done by Welch’s T-test.

### Combined inhibition of both Adenosine receptors A2A and A2B is necessary to prevent the effect of adenosine on glucose and fatty acid metabolism

Caffeine is a pan-adenosine receptor antagonist that has a half-life of about one hour in mice following i.p. injection [31]. We sought to investigate whether administration of caffeine before an adenosine challenge would be sufficient to block the adenosine effects on glucose and NEFA. Refed WT mice were given an i.p. bolus of caffeine (25 mg/kg) and, after a 30-minute equilibration period, were given a bolus of adenosine (100 mg/kg). Mice injected with caffeine had reduced glucose (Fig. 3A) and NEFA excursions (Fig. 3B) compared to mice given adenosine alone. To test the necessity of the A2A and A2B receptors in promoting this effect, we used the A2A-selective antagonist SCH-58261 and the A2B-selective antagonist ATL-844 before an adenosine challenge. Refed mice were injected with either SCH-58261 (3 mg/kg) or ATL-844 (3 mg/kg). After a 30-minute equilibration period, mice were given a bolus of adenosine (100 mg/kg). Neither SCH-58261 nor ATL-844 treatment alone was sufficient to block the adenosine effect on glucose (Fig. 3 C) or NEFA (Fig. 3 D). However, when mice were given a concomitant injection SCH-58261 and ATL-844, both the adenosine-mediated glucose (Fig. 3 E) and NEFA (Fig. 3 F) excursions were abolished. Fasted mice did not show a significant difference in glucose or NEFA excursion between drug and vehicle-treated groups. Refed female mice given a concomitant injection of SCH-58261 and ATL-844 before an adenosine challenge also showed significantly lower glucose and NEFA responses (Sup. Fig. 1 G, H), demonstrating that this is not a sex-specific phenomenon.

**Figure 3:**
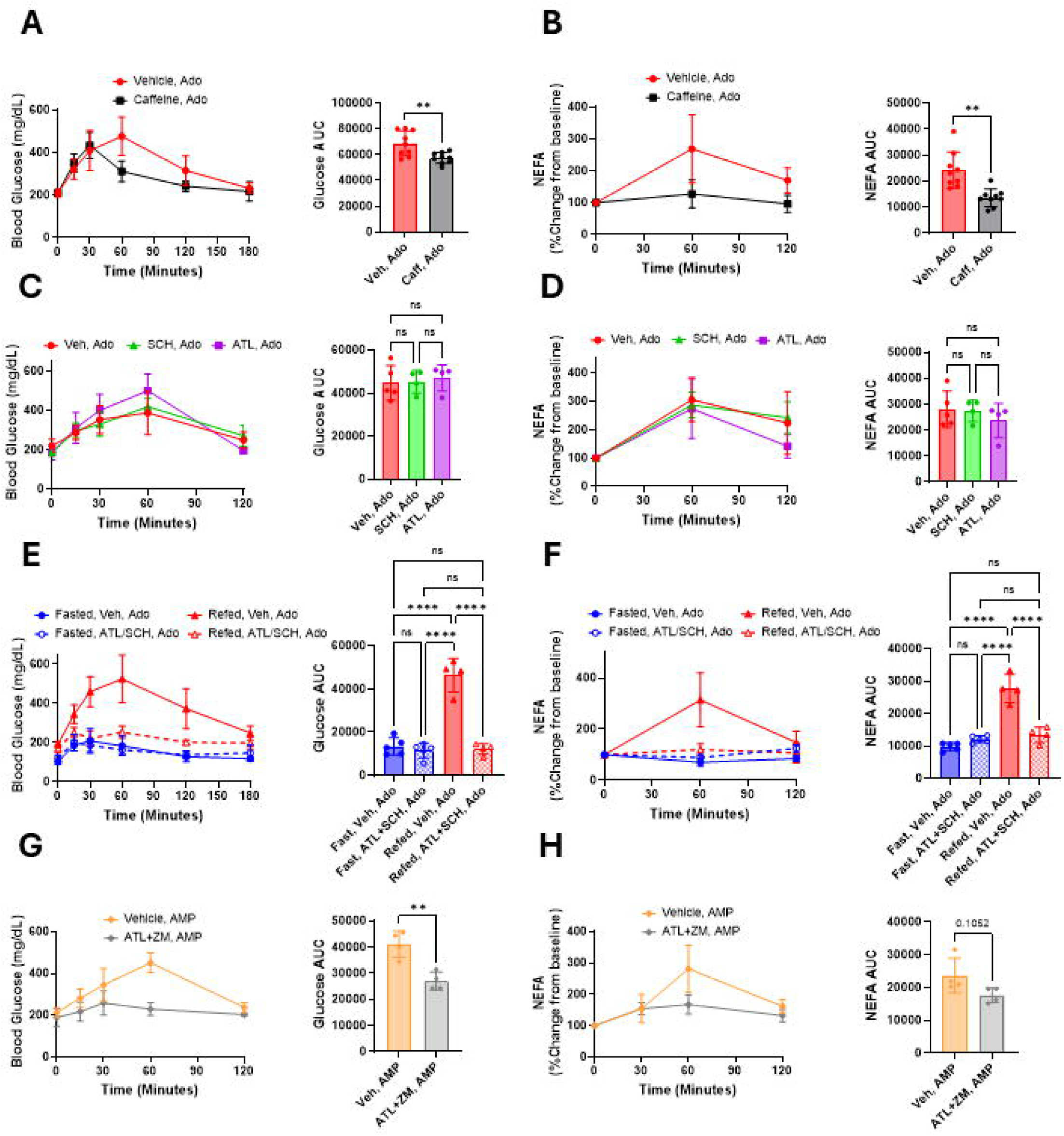
A2A and A2B are individually sufficient, but collectively necessary for postprandial adenosine-induced glucose and NEFA excursion. WT mice were fasted and refed as previously mentioned. At t = -30 minutes, refed mice were i.p. injected with either vehicle or caffeine (25 mg/kg). At t = 0 minutes, all mice received a bolus of adenosine (100 mg/kg). **(A)** Blood glucose and **(B)** serum NEFA were tracked over 2 hours. WT refed mice were injected at t = -30 minutes with an i.p. bolus of vehicle (red), SCH-58261 (3 mg/kg, green), or ATL-844 (3 mg/kg, violet). At t = 0, all mice received an i.p. bolus of adenosine (100 mg/kg). **(C)** Changes in blood glucose levels and **(D)** serum NEFA were recorded over 2 hours. WT fasted and refed mice were i.p. injected with either vehicle or a cocktail of both ATL-844 and SCH-58261 (ATL+SCH, 3 mg/kg each) at t = -30 minutes. At t = 0, all mice were given an i.p. bolus of adenosine (100 mg/kg). **(E)** Blood glucose levels and (**F)** serum NEFA of fasted/vehicle/adenosine (dark blue), fasted/ ATL+SCH/adenosine (spotted, light blue), refed/vehicle/adenosine (dark red), and refed/ATL+SCH/adenosine (spotted, light red) mice plotted over the course of 3 hours. WT, refed mice were i.p. injected with vehicle + 5’ AMP (150 mg/kg) or a cocktail of 5’ AMP (150 mg/kg) + ATL-844 + SCH-58261 (3 mg/kg, each). **(G)** Blood glucose and **(H)** serum NEFA was recorded over a two-hour period. Error bars represent SD. *p < 0.05, **p < 0.01, ***p < 0.001, ****p < 0.0001. All statistical analyses were done by Welch’s T-test (A-B, G-H), one-way ANOVA (C-D), or two-way-ANOVA (E-F).

Administration of 5’ AMP can also promote acute glucose excursion in mice [32,33]. In agreement with previous reports, giving mice an i.p. bolus of 5’ AMP at 150 mg/kg was sufficient to promote a large increase in blood glucose (Fig. 3 G). Interestingly, we also found that there was a concomitant increase in non-esterified fatty acids (NEFA) indicating that adipocyte lipolysis was being stimulated (Fig. 3 H). Refed mice given a simultaneous dose of SCH-58261 and ATL-844 before an 5’ AMP bolus had a diminished glucose and NEFA excursion compared with mice treated with 5’ AMP alone (Fig 3 G, H), while treatment with just the A2A antagonist ZM- 241385 (Sup. Fig. 2 A, B) or ATL-844 (Sup. Fig. 2 C, D) were insufficient to affect glucose or NEFA.

To complement these pharmacological studies, we crossed previously described single floxed alleles of *Adora2A* and *Adora2b* to generate a global double knockout line. Mice were bred to contain homozygous floxed alleles of both *Adora2a* and *Adora2b* genes under the ubiquitin-Cre-ERT2 and were dubbed “UBC-A2A+B”. At eight weeks both Cre+ and Cre- mice were given tamoxifen at 75 mg/kg for five days, followed by a two-week washout period (Sup. Fig. 3 A, B) UBC-A2A+B WT (Cre-) and UBC-A2A+B DKO (Cre+) mice were then given an adenosine challenge (100 mg/kg) under fasted and refed conditions. We found that under fasted conditions, UBC-A2A+B DKO mice had a slight but significantly lower adenosine-mediated glucose excursion compared with UBC-A2A+B WT (Fig. 4 A), while NEFA levels did not change in either group. Under refed conditions, UBC-A2A+B DKO mice showed a significant reduction in both adenosine-mediated glucose and NEFA release compared with UBC-A2A+B WT mice (Fig. 4 C, D). Thus, both genetic ablation and pharmacological inhibition demonstrate that simultaneous blockade of A2A and A2B receptors is required to suppress adenosine’s metabolic effects.

**Figure 4:**
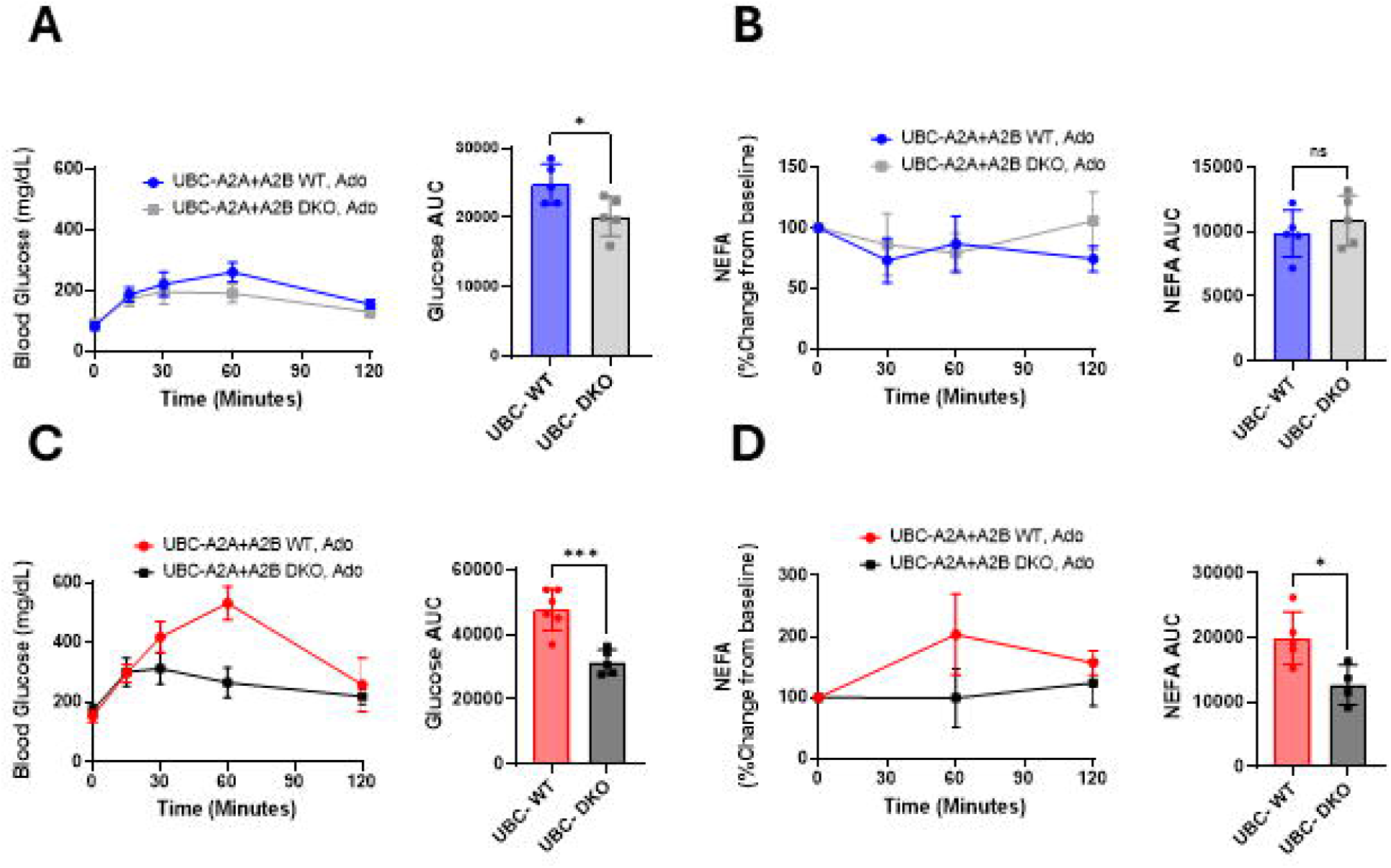
Global genetic ablation of A2A and A2B receptors diminishes the postprandial adenosine effect on glucose and NFEA. Fasted UBC-A2A+B WT and UBC-A2A+B DKO mice were given an i.p. bolus of adenosine (100 mg/kg). **(A)** Blood glucose and **(B)** serum NEFA were recorded over a two-hour period. Refed UBC-A2A+B WT and UBC-A2A+B DKO mice were given an i.p. bolus of adenosine (100 mg/kg) and **(C) b**lood glucose and **(D)** serum NEFA were recorded over a two-hour period. N’s for each group range from 4-6. Error bars represent SD. *p < 0.05, **p < 0.01, ***p < 0.001, ****p < 0.0001. All statistical analyses done by Welch’s T-test.

### Adenosine promotes glycogenolysis in the liver and lipolysis in the adipose tissue

We next sought to determine which tissues were responsible for mediating the adenosine effects on glucose and NEFA. Liver and adipose tissue are the master regulators of glucose and lipid homeostasis in the body, respectively. We sought to determine if there were catabolic signaling changes in liver and WAT after an adenosine challenge. Refed WT mice were given an i.p. bolus of vehicle or adenosine (100 mg/kg) and tissues were then harvested 30 minutes later. Adenosine significantly increased levels of phosphorylated CREB in the livers and gonadal white adipose tissue compared to vehicle controls (Fig. 5 A, B), suggesting that catabolic signaling was indeed activated in these tissues in response to adenosine. The liver is able to rapidly increase blood glucose levels in response to hormonal signals via glycogenolysis [34]. We next investigated whether adenosine administration stimulates glycogenolysis by measuring hepatic glycogen levels after an adenosine challenge. Fasted WT mice were given an i.p. bolus of vehicle, and refed WT mice were given either vehicle or adenosine (100 mg/kg). After one hour, all mice were sacrificed and their livers were harvested. Measurement of hepatic glycogen revealed that adenosine treatment significantly reduced liver glycogen content in refed mice compared with vehicle (Fig. 5C), although levels remained higher than in overnight-fasted mice. We next wanted to investigate whether adenosine affects insulin levels, which would contribute to elevated circulating glucose levels. Refed WT mice were given an i.p. bolus of adenosine (100 mg/kg), BAY-60-6583 (BAY) (0.2 mg/kg), or a concomitant dose of SCH-58261 + ATL-844 (3 mg/kg, each) followed by a bolus of adenosine 30 minutes later. Serum was collected before and 60 minutes after drug treatment. For SCH+ATL-treated mice, t = 0 serum was taken 30 minutes after ATL+SCH injection, but before adenosine injection. Both adenosine and BAY treatment was sufficient to significantly lower circulating insulin levels one-hour after injection, and pre-treatment with SCH+ATL was sufficient to prevent adenosine-induced decrease in insulin (Fig 5 D).

**Figure 5:**
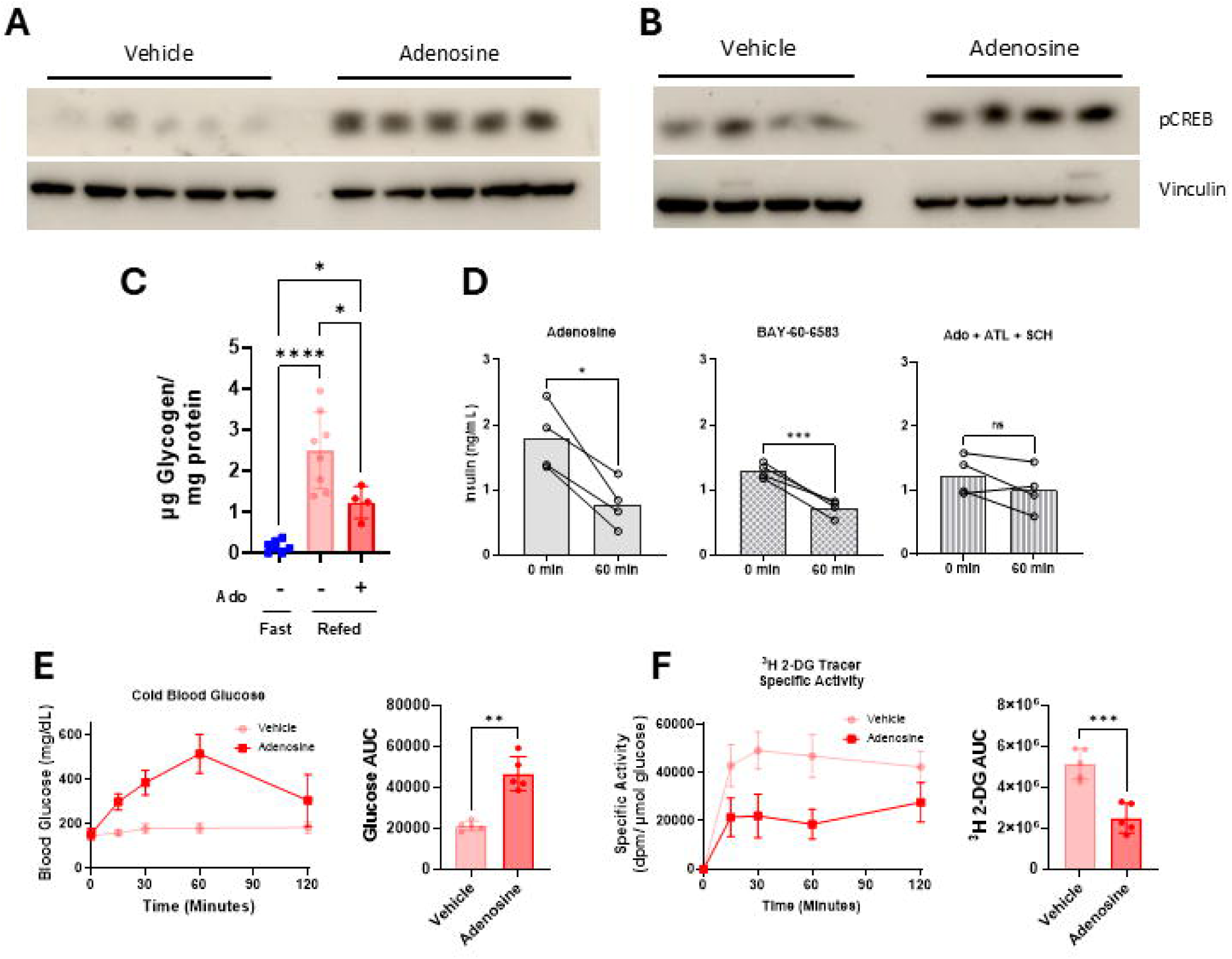
Adenosine stimulates hepatic glucose production and adipocyte lipolysis. WT, refed mice were i.p. injected with either vehicle or adenosine (100 mg/kg). After 30 minutes, animals were sacrificed. Levels of phosphorylated CREB were measured in **(A)** liver and **(B)** gonadal white adipose tissue (GWAT). **(C)** Liver glycogen levels extracted from WT mice that were either: fasted, refed, or refed and treated with an i.p. bolus of adenosine for 1 hour. **(D)** Serum insulin levels measured in refed WT mice before and 1 hour after i.p. injection of adenosine (100 mg/kg), BAY (0.2 mg/kg), or adenosine after 30-minute pretreatment with ATL-844 and SCH-58261 (3 mg/kg, each). WT mice were injected with 10 µCi of ^3^H-2-DG tracer +/-adenosine (100 mg/kg). Blood glucose and serum were collected over a two-hour period, then the animals were sacrificed. **(E)** Levels of blood glucose (cold) over the two-hour period. **(F)** Specific activity of ^3^H-2-DG tracer in serum over the two-hour period. N’s for each group range from 4-8. Error bars represent SD. *p < 0.05, **p < 0.01, ***p < 0.001, ****p < 0.0001. All statical analyses done by one-way ANOVA (C) or Welch’s T-test (D-F).

To study more directly how adenosine affects glucose homeostasis, we investigated glucose appearance in the blood by dilution of a radiolabeled glucose tracer. At t = 0, refed mice were given a dose of [^3^H]- 2-DG (10 µCi) with or without adenosine (100 mg/kg). Serum was collected over a two-hour period, after which their tissues were harvested. Consistent with our previous observations, cold glucose levels increased drastically in the adenosine-treated group (Fig. 5 E). The amount of circulating tracer was normalized to circulating cold glucose levels to generate specific activity (disintegrations per minute (dpm) / µM circulating glucose). We found that adenosine-treated mice had significantly lower serum specific activity compared with vehicle-treated mice (Fig. 5 F) demonstrating a rapid increase in cold glucose. Together these data indicate that administration of adenosine to refed mice activates catabolic signaling in liver and adipose tissue, promotes liver glycogenolysis and adipose tissue lipolysis, and greatly increases the apparent rate of glucose appearance. These catabolic processes are likely potentiated by the concomitant suppression of circulating insulin.

### Cell autonomous A2A and A2B signaling in the adipose tissue and liver are not necessary for adenosine’s effects on NEFA and glucose, respectively

Given that both A2A and A2B couple Gs, we hypothesized that adenosine action on A2A and A2B receptors on the liver and adipose tissue was driving cAMP/PKA activity and driving hepatic glucose production and adipocyte lipolysis. We crossed A2A and A2B homozygous double-floxed mice with the adipoQ-Cre-ERT2 line to generate adipocyte-specific Cre A2A + A2B double floxed mice (F-A2A+B). We anticipated that deletion of both A2A and A2B in adipose tissue would eliminate the adenosine-mediated effects on lipolysis and circulating NEFA. FA2A+B, Cre- (WT) and FA2A+B, Cre+ (DKO) mice were treated with tamoxifen and given a washout period as previously described. Surprisingly, when refed FA2A+B DKO mice were given an i.p. bolus of the A2B agonist BAY-60-6583 (BAY) (0.2 mg/kg), there was no change in the magnitude in NEFA excursion compared to F-A2A+B WT mice (Sup. Fig. 4 B) and glucose was also unaffected (Sup. Fig. 4 A).

In parallel to developing our F-A2A+B mice, we also produced A2A and A2B liver double knockout mice. A2A and A2B double floxed mice were given intravenous tail vein injections of either AAV8-TBG-Cre or AAV8-TBG-GFP to generate A2A + A2B liver-double knockout (AAV-Cre) and WT control (AAV-GFP) mice. BAY did not elicit a significant difference in glucose excursion between WT and liver DKO mice (Sup Fig. 4 C), nor was there a difference in NEFA excursion (Sup Fig. 4 D). These data indicate that A2A and A2B signaling in adipose tissue and the liver are not necessary for the adenosine effect. Taken together with the previous data showing the necessity of A2A and A2B receptors in mediating the adenosine effect, we surmised that A2A and A2B signaling must be acting in a cell non-autonomous way on the adipose tissue and liver.

### Adenosine mediates glucose and NEFA modulation via sympathetic signaling

We considered what other signaling pathways might contribute to the metabolic effects seen with acute adenosine administration. The rate at which both NEFA and glucose increase after an adenosine bolus suggested that there must be an abrupt activation of a secondary signaling pathway. Activation of PKA/CREB signaling in liver and adipose tissue is consistent with adrenergic signaling [35] and adenosine has been previously linked to modulating catecholamine release at both pre- and postganglionic neurons [36–39] and the adrenal medulla [40,41].

To investigate whether adenosine was affecting glucose and NEFA levels by increasing sympathetic activation, we treated mice with hexamethonium before and adenosine challenge. Hexamethonium is a nicotinic acetylcholine receptor (nAChR) antagonist that functions to inhibit autonomic signaling. Refed mice were given an i.p. bolus of vehicle or hexamethonium (30 mg/kg) at t = -30. At t = 0, all mice received an additional i.p. bolus of adenosine (100 mg/kg). Mice treated with hexamethonium had significantly lower glucose and NEFA release in response to adenosine compared with vehicle controls (Fig. 6 A-B). Next, we sought to pharmacologically inhibit the sympathetic pathway by blocking both α and β adrenergic receptors. Refed WT mice received either an i.p. bolus of adenosine (100 mg/kg) alone, or a bolus of adenosine (100 mg/kg) along with a concomitant dose of the α -adrenergic receptor antagonist phentolamine (10 mg/kg) and the β-adrenergic receptor antagonist propranolol (10 mg/kg). Similar to the mice treated with hexamethonium, mice that were treated with the α and β adrenergic receptor antagonists showed a marked reduction in the ability of adenosine to increase blood glucose and NEFA (Fig. 6 C-D). We conclude from these data that sympathetic signaling is required for mediating adenosine-stimulated effects on blood glucose and serum NEFA.

**Figure 6:**
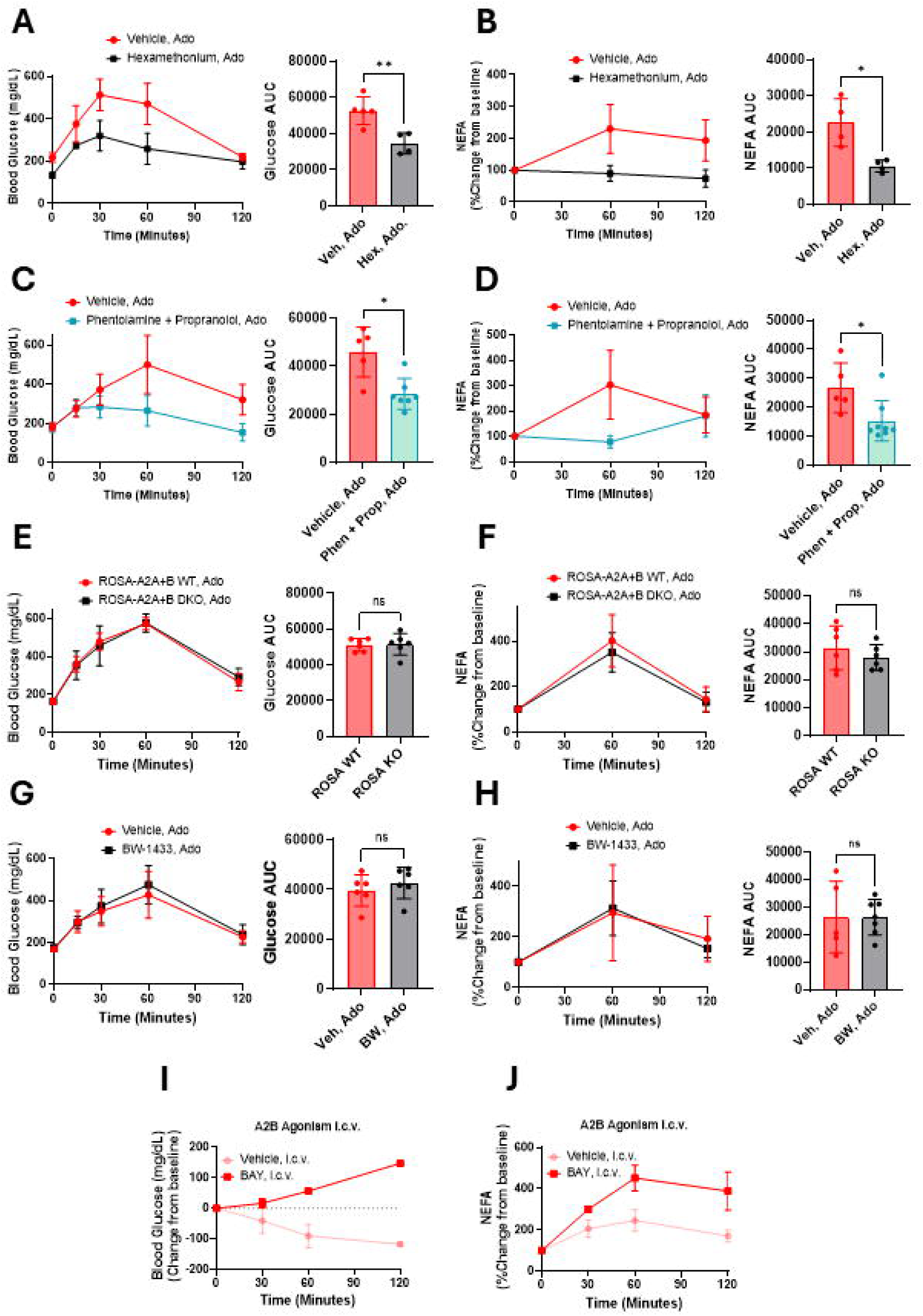
Adenosine- mediated glucose and NEFA excursion requires sympathetic signaling. At t = -30 minutes, a group of refed WT mice were i.p. injected with either vehicle or a combination of phentolamine and propranolol (10 mg/kg each) (N = 5). At t = 0 minutes, all mice received an i.p. bolus of adenosine (Ado) (100 mg/kg). **(A)** Blood glucose and **(B)** serum NEFA was recorded over 2 hours. In a separate experiment a group of refed WT mice were i.p. injected with vehicle or the autonomic signaling inhibitor hexamethonium (Hex) (30 mg/kg) at t = -30 minutes. At t = 0 minutes, all mice received an i.p. bolus of adenosine (100 mg/kg). **(C)** Blood glucose and **(D)** Serum NEFA was recorded over 2 hours. ROSA26-A2A+B were refed and given adenosine and **(E)** blood glucose and **(F)** and serum NEFA were recorded over a two-hour period. WT refed mice were i.p. injected with vehicle or BW-1433 (10 mg/kg) at t = -60. AT t = 0, all mice received adenosine (100 mg/kg). **(G)** Blood glucose and **(H)** serum NEFA were recorded over a two-hour period. Refed WT mice received an intracerebroventricular (ICV) injection of vehicle or BAY-60-6583 (0.02 mg/kg). **(I)** Change in blood glucose from baseline and **(J)** serum NEFA was tracked over a two-hour period. N’s for each group range from 4-8. Error bars represent SD. *p < 0.05, **p < 0.01, ***p < 0.001, ****p < 0.0001. All statistical analyses done by Welch’s T-test.

### A2A and A2B signaling in peripheral neurons is unnecessary for the adenosine effect

Considering that adrenergic receptors seemed to be required for mediating the adenosine effect, and that this effect requires sympathetic signaling, we hypothesized that double knockout of A2A and A2B in neurons would be sufficient to block the adenosine effects on glucose and NEFA. In order to discern whether adenosine was acting in the central or peripheral nervous system, we sought to selectively delete A2A and A2B in peripheral neurons. To do this, we implemented an AAV9-PHP.S virus expressing Cre recombinase under control of the hSYN1 promoter. The PHP.S capsid has been shown previously to specifically target peripheral neurons, but fails to cross the blood brain barrier [25]. Moreover, the hSYN1 promoter is uniquely expressed in neurons, further adding to the specificity of the virus to peripheral neurons. A2A + A2B double floxed mice were given AAV9-PHP.S-hSYN1-Cre-GFP or AAV9-PHP.S-CAG-TdTomato as an infection control. Mice were injected with 1×10^12^ GC/ mouse via tail vein injection and were given a three-week washout period. Refed AAV9-tdTomato (WT) and AAV9-Cre (A2A+B DKO) were then challenged with an i.p. injection of 100 mg/kg adenosine. AAV-Cre mice had a slight, but significant decrease in blood glucose excursion (Sup Fig. 5 A), but there was no difference in NEFA between AAV-Cre and AAV-GFP groups (Sup Fig. 5 B). Next, we challenged refed AAV9-tdTomato and AAV9-Cre refed mice with an i.p. bolus of ATL-313 (0.2 mg/kg) or BAY (0.2 mg/kg). In both instances, there was no significant difference in the magnitude of glucose or NEFA excursion between AAV9-tdTomato and AAV-9Cre groups (Supp Fig 5 C-D). Successful infection of peripheral neurons was evident by confirming eGFP-expression neurons of the superior cervical ganglion (SCG). (Sup Fig. 5 G). Despite a subtle reduction in blood glucose excursion when challenged with adenosine, knockout of A2A and A2B in peripheral neurons failed to robustly block the adenosine effects to the same degree seen with dual pharmacological inhibitors ATL+SCH treatment or the UBC-A2A+B DKO mice. We conclude from these data that A2A and A2B signaling in peripheral neurons has a very minor role in facilitating the adenosine effect on glucose metabolism and has no effect on NEFA metabolism.

To complement our peripheral neuron KO mice, we crossed A2A + A2B double floxed mice with the ROSA26-Cre-ERT2 line. These mice have a tamoxifen-inducible Cre that has very weak recombination activity in the brain but otherwise successfully recombines in all other peripheral tissues when tamoxifen is administered i.p. [42,43]. The resulting ROSA-A2A+B Cre + mice have WT expression of A2A and A2B in the brain but are globally deleted in all peripheral tissues (Sup. Fig. 6 C, D). When ROSA26-A2A+B DKO mice were challenged with an adenosine bolus, there was no change in the level of glucose or NEFA excursion compared with ROSA26-A2A+B WT mice. Finally, we sought to pharmacologically inhibit peripheral adenosine signaling and examine the effects on the adenosine mediated glucose and NEFA excursion. BW-1433 is a pan-adenosine antagonist with high affinity to each receptor (K_d_: A1 = 140 nM, A2A = 190 nM, A2B = 60 nM, A3 = 30 nM) [44] that does not cross the blood brain barrier [45]. WT refed mice were given an i.p. bolus of vehicle or BW-1433 (10 mg/kg) then, after a 60-minute equilibration period, were given a bolus of adenosine (100 mg/kg). There was no difference in the magnitude of blood glucose excursion (Fig 6 G) or NEFA excursion (Fig 6 H) between mice treated with BW-1433 and adenosine versus mice treated with adenosine alone. We next sought to investigate if direct injection of an adenosine agonist into the central nervous system would be sufficient to induce glucose and NEFA release. Refed WT mice received intracerebroventricular (i.c.v.) injection of vehicle or BAY-60-6583 (0.02 mg/kg). Mice that received an i.c.v. injection of BAY experienced a significantly higher increase in both glucose and NEFA compared to vehicle-treated controls (Fig. 6 I, J). Importantly, refed WT mice injected with the same dose of BAY i.p. did not display any significant glucose or NEFA excursion compared to vehicle controls (Sup. Fig. 6 H, I). These genetic and pharmacological data strongly suggest that central adenosine signaling is required for the metabolic effects of adenosine.

### A2A and A2B double knockout in neurons is sufficient to block the adenosine effect

Considering that sympathetic signaling appears to be required for the adenosine effect, while deletion of A2A and A2B in peripheral neurons is insufficient to block the adenosine effect, we hypothesized that double knockout of A2A and A2B in all neurons (both peripheral and central) would be sufficient to block the adenosine effects on glucose and NEFA. To generate a pan-neuron-specific A2A and A2B double knockout line, we crossed A2A and A2B double floxed mice with the Baf53b-Cre mouse line, to direct Cre expression via the *Actl6b* promoter in all neurons (both central and peripheral) (Sup. Fig. 6 A, B) [46]. Similar to the UBC-A2A+B DKO mice, we found that fasted Baf-A2A+B DKO (Cre -) mice showed a slight but significantly lower glucose excursion in response to adenosine compared with Baf-A2A+B WT controls (Fig. 7 A). Both Baf-A2A WT and DKO mice had no change in serum NEFA under fasted conditions (Fig 7 B). However, refed Baf-A2A+B DKO mice showed a significantly lower excursion of glucose and NEFA compared with WT controls (Fig. 7 C-D). We also tested the effect of individual A2A and A2B agonism in refed Baf-A2A+B WT and DKO mice. Baf-A2A+B DKO mice showed a significantly lower glucose and NEFA excursion compared to WT controls when challenged with both ATL-313 (Fig. 7 E, F) and BAY-60-6583 (Fig. 7 G, H). Collectively, these data implicate A2A and A2B signaling in central neurons as being necessary for the adenosine-mediated effect on blood glucose and NEFA.

**Figure 7:**
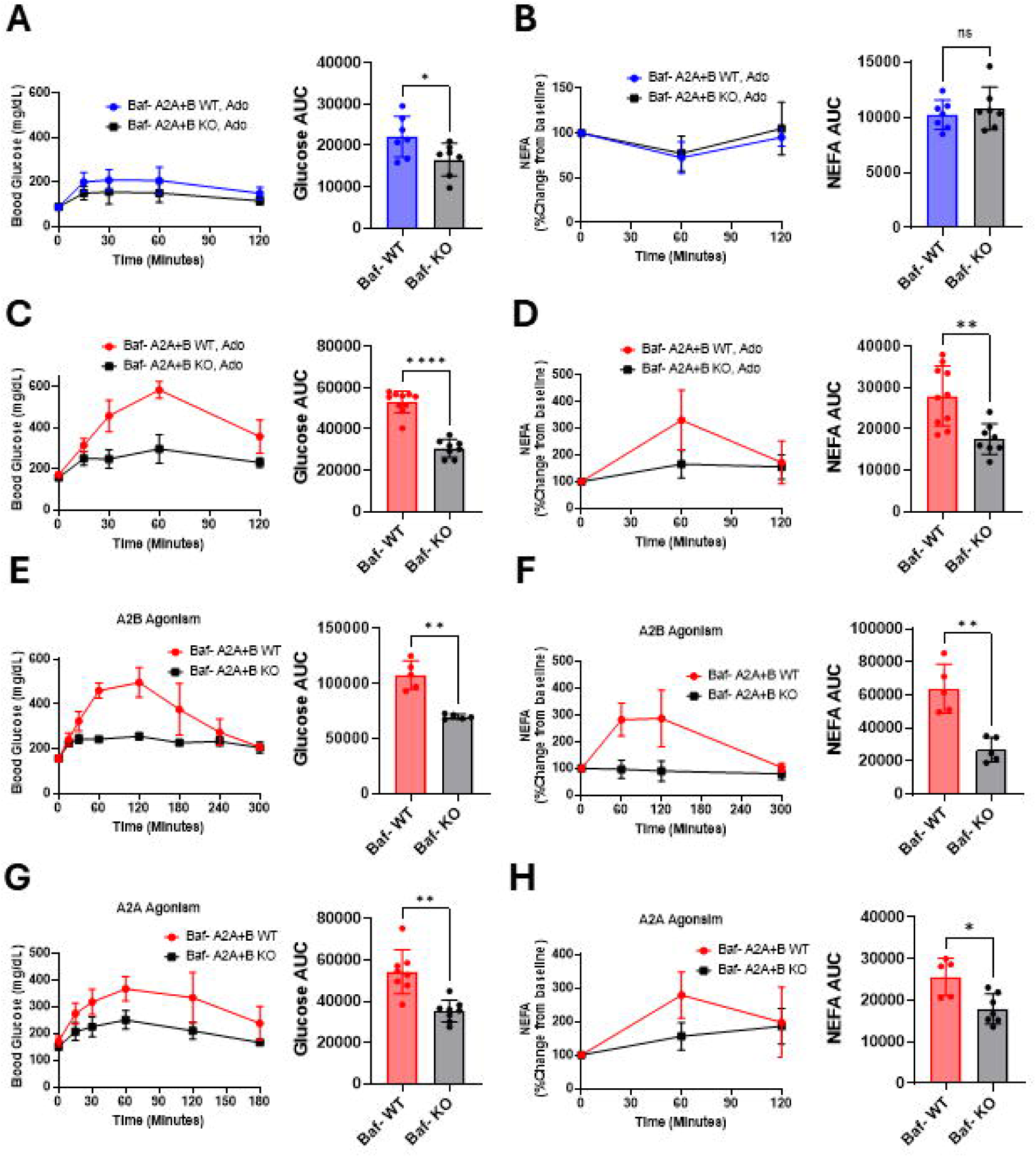
Baf53b-A2A+A2B double knockout mice have a diminished metabolic response to an adenosine challenge. A neuron-specific A2A+A2B double knockout mouse line was generated using the neuron-specific Baf53b promoter Baf53b-A2A+B WT and DKO mice were injected with an i.p. bolus of adenosine (100 mg/kg) and **(A)** Blood glucose and **(B)** serum NEFA levels were measured over a 2-hour period. This experiment was then repeated in refed Baf53b-A2A+B WT (N= 5) and KO (N = 5) mice and **(C)** Blood glucose and **(D)** serum NEFA were also measured over a 2-hour period. In a separate experiment, refed Baf53b-A2A+B WT (N = 5) and KO (N = 5) mice were challenged with an i.p. bolus of the A2B agonist BAY 65-6083 (0.2 mg/kg) and **(E)** blood glucose and **(F)** serum NEFA were recorded over 5-hour period. Refed Baf53b-A2A+B WT (N = 8) and KO (N = 8) were challenged with an i.p. bolus of the A2A agonist ADO-313 (0.2 mg/kg) and **(G)** blood glucose and **(H)** serum NEFA were measured over a three-hour period. N’s for each group range from 5-10. Error bars represent SD. *p < 0.05, **p < 0.01, ***p < 0.001, ****p < 0.0001. All statistical analyses were done by Welch’s T-test.

## DISCUSSION

This study shows that administration of a single bolus of adenosine is sufficient to drive an acute and profound increase in blood glucose and NEFA in the postprandial state. We demonstrate that this effect can be blocked by simultaneous administration of selective A2A and A2B antagonists. Importantly, we observed that each inhibitor was insufficient by itself to affect the adenosine effects on metabolism, and that these effects were only abolished when both A2A and A2B receptors were inhibited simultaneously. As an orthogonal approach to pharmacology, we utilized genetic ablation of A2A and A2B in the whole body using a UBC-A2A+B mouse line. Similar to the double antagonist experiment, the adenosine metabolic response was completely abolished in UCB-A2A+B DKO mice. This suggests there is functional redundancy between these receptors, such that signaling through either one alone is sufficient to elicit the metabolic effects. We conclude that stimulation of either receptor is sufficient to promote glucose and NEFA release, but both receptors must be simultaneously inhibited to abolish adenosine action. We summarize this as both A2A and A2B receptors are individually sufficient, but collectively necessary for facilitating the metabolic effects of adenosine.

Similar to adenosine, a bolus of 5’ AMP elicited an acute, robust increase in both glucose and NEFA. This agrees with previous reports where 5’ AMP rapidly increases blood glucose [32,33]. Interestingly we show here that these 5’ AMP-mediated changes in blood glucose and NEFA are blocked when the A2A and A2B antagonists SCH-58261 and ATL-844 are injected 30 minutes prior. Unlike ATP and ADP, 5’ AMP is incapable of activating P2Y and P2X receptors, precluding their involvement in this process [47]. Additionally, 5’ AMP is generally thought to be unable to activate adenosine receptors, unless it is first converted to adenosine by CD73 [48]. Our findings suggest that the glucose and NEFA effects seen with a 5’ AMP bolus are being mediated by A2A and A2B receptors after 5’ AMP is converted to adenosine in the extracellular space.

Compared to fasted mice, postprandial mice undergo a much more profound and sustained release of glucose in response to adenosine. Intuitively, the dichotomy of glucose responses between postprandial and fasted mice might be explained by substrate availability. Adenosine administration in refed mice promotes activation of the PKA/ CREB pathway within 30 minutes, and significant reduction in liver glycogen as soon as one-hour after treatment, which coincides with the peak in circulating glucose. Moreover, the increased rate of glucose appearance seen in refed mice during an adenosine challenge is consistent with glucose release from hepatic glycogen. This contrasts with fasted mice, whose glycogen levels are nearly undetectable at the start of the experiment, and adenosine only elicits a minor glucose release.

Superficially, the magnitude of glucose release stimulated by adenosine is then proportional to the level of glycogen in the liver at the time of the experiment. However, glucagon, a hormone that directly stimulates hepatic glucose production, promotes a similar level of glucose production as adenosine under fasted conditions, but under postprandial conditions the effect of glucagon on blood glucose is dwarfed by that of adenosine. The difference in these postprandial glucose responses can likely be attributed to adenosine’s ability to suppress circulating insulin levels. We show that blood insulin levels are significantly lowered in refed mice one hour after administration of adenosine, and that this effect can be blocked by pretreatment with A2A and A2B antagonists. The A2B agonist BAY-60-6583 was also sufficient to significantly lower blood insulin after one hour, indicating this effect is due to adenosine receptor activation, and not an off-target effect of adenosine or its metabolites. This postprandial drop in insulin would be expected to potentiate catabolic signaling in liver and adipose tissue and promote liberation of stored glucose and NEFA. Glucagon is incapable of suppressing blood insulin levels, and in fact has been shown to increase insulin release when given at high doses [49]. It remains unclear whether the effects of adenosine on insulin are due to a cell autonomous or cell non-autonomous effect on pancreatic beta cells. Baf-A2A+B DKO mice do not show a suppression of insulin after an adenosine challenge (Sup. Fig 7 A-B), implicating central adenosine signaling in this process. However, more work needs to be done in this area to clarify the exact mechanism.

Similar to glucose, adenosine substantially increases circulating NEFA levels in the postprandial state. In contrast, adenosine has little effect on lipolysis in the fasted state. This difference can likely be attributed to cell-autonomous adenosine signaling on adipose tissue. Adenosine has long been known to inhibit lipolysis in isolated adipocytes via the A1 receptor [50]. We have previously reported that this effect is potentiated in mice under fasted conditions due to transcriptional upregulation of the A1 receptor in adipose tissue [27]. A1 protein and mRNA levels are high in adipose tissue under fasted conditions but decrease abruptly after refeeding. Accordingly, isoproterenol-induced lipolysis was significantly reduced by A1 agonism in the fasted state, but not refed state. In the present study these findings suggest that in the fasted state, elevated adipose A1 expression allows adenosine to oppose lipolysis despite concurrent adenosine-mediated catecholamine release, leading to no net change in serum NEFA levels. However, under refed conditions, A1 levels in adipose tissue decrease, circulating insulin levels decrease, and catecholamine-induced lipolysis can proceed unfettered by adenosine suppression of lipolysis in adipocytes.

Global double deletion of A2A and A2B substantially lowers the glucose and NEFA response during a refed adenosine challenge. These genetic data complemented the results from simultaneous pharmacological antagonism of A2A and A2B signaling, but left unanswered the question of which tissues were required for this response. Activation of PKA/ CREB signaling in the liver and adipose tissue occurred in parallel with adenosine-induced changes in glucose and fatty acid metabolism, potentially suggesting direct adenosine activation of the Gs-coupled A2A and A2B receptors on these tissues could be mediating these metabolic changes. However, deletion of both adenosine receptors in adipose tissue did not dampen the adenosine-induced increase in glucose and NEFA in the refed state. Similarly, double deletion of A2A and A2B in the liver by means of a AAV8-TBG-Cre virus showed no impairment of adenosine-mediated glucose or NEFA excursion under refed conditions. These data suggested that adenosine signaling acts on adipose and liver tissue in a cell non-autonomous manner.

Since deleting the A2 receptors in adipose tissue and liver had no effect, we speculated that adenosine’s actions were instead mediated by autonomic circuits, which are well known to stimulate adipose tissue lipolysis and hepatic glucose production. Hexamethonium markedly reduced the adenosine-induced increases in glucose and NEFA, consistent with autonomic activity being required. Simultaneous α- and β-adrenergic blockade indicates that sympathetic signaling is a necessary component of adenosine-mediated activation of adipose tissue and liver. These experiments do not address whether the released catecholamines originated from sympathetic neurons directly innervating to adipose tissue and liver, or from sympathoadrenal epinephrine release, or whether the magnitude of these responses differs between fasted and refed states.

Given the requirement for autonomic signaling, we were curious as to whether adenosine was acting on A2A and A2B receptors directly on peripheral or central neurons. Afferent neurons in the carotid body have been shown to be responsive to endogenous adenosine [51], and are also known to be able to stimulate sympathetic signaling in response to chemical stimuli (e.g. low O_2_ during hypoxia, or high CO_2_ during acidosis) [52]. In addition, adenosine has been reported to modulate catecholamine release via presynaptic adenosine receptors, with A1-mediated inhibition [37,38] and A2-mediated stimulation [36,39] acting in opposing directions to tune sympathetic output. To assess the involvement of adenosine receptors in peripheral neurons, we selectively deleted A2A and A2B in peripheral neurons using the AAV9-PHP.S-Cre virus. Chan and colleagues showed this viral serotype to be highly penetrant into peripheral neurons, but unable to cross the blood brain barrier and infect central neurons [25]. Ganglionic eGFP expression confirmed high infectivity of the Cre expressing virus in peripheral neurons, but these animals showed no impairment in NEFA, and only a minimal decrease in glucose release, when challenged with adenosine or A2A/A2B agonists.

In parallel to our virus treatment scheme, we also developed an A2A and A2B double floxed ROSA26-Cre-ERT2 mouse line. Several studies have demonstrated that when moderate doses of tamoxifen are administered intraperitoneally, these mice undergo Cre-mediated recombination in peripheral tissues but not in the brain [42,43]This has been ascribed to low ROSA26 promoter activity in neural tissue. ROSA26-A2A+A2B DKO mice showed an equal adenosine response to WT controls, further supporting the notion that peripheral adenosine signaling is not necessary for the adenosine effects on metabolism. Finally, to investigate the pharmacological effects of inhibiting peripheral adenosine receptors, we used the peripherally restricted adenosine antagonist, BW-1433. BW-1433 has a much higher affinity for adenosine receptors than caffeine (BW-1433 K_d_: A1 = 140 nM, A2A = 190 nM, A2B = 60 nM, and A3 = 30 nM [44] versus caffeine K_d_: A1 = 12 µM, A2A = 2.4 µM, A2B = 13 µM, and A3 = 80 µM [53]). However, unlike caffeine, the carboxylic acid BW-1433 cannot penetrate the blood brain barrier [44].

Administration of BW-1433 at a saturating dose before an adenosine challenge had no discernable impact on adenosine-mediated glucose and NEFA excursion, whereas caffeine was sufficient to greatly reduce this effect. Finally, Baf-A2A+B DKO mice given adenosine, an A2A agonist (ADO-313), or A2B agonist (BAY-60-6583) all had a significantly lower glucose and NEFA response compared to WT controls. These data support a role for A2A and A2B signaling in central neurons, not peripheral neurons, for adenosine’s metabolic effects.

Although our work indicates a requirement of adenosine signaling in the CNS, it remains unclear which region of the brain is required. The hypothalamus would be a reasonable candidate as it has well described roles in blood glucose homeostasis, adipocyte lipolysis, and energy expenditure [54–56] Indeed, many of these effects are mediated by modulating autonomic signaling. Candidate regions are hypothalamic nuclei that integrate metabolic signals and project to pre-autonomic neurons, especially the paraventricular nucleus (PVN) and nearby hypothalamic regions such as the arcuate nucleus (ARC) and lateral hypothalamus (LH). These regions may express adenosine receptors on neurons and are known to drive changes in sympathetic and parasympathetic outflow to liver, pancreas, and adipose tissue, thereby altering glucose and NEFA levels. It is important to note that while A2A and A2B are both individually sufficient and genetic data supports neuronal expression, it is unclear whether these receptors are expressed on the same neurons. In fact, it is quite possible that different regions of the brain display differential receptor expression. The substantial difference in affinity for adenosine between A2A and A2B (A2A K_i_ = 310 nM vs A2B K_i_ = 15,000 nM in humans) [30] suggests that they could be mediating different signals or displaying a biphasic response to different levels of adenosine whereby A2A is activated under low levels and A2B gets activated under high levels. Finally, it is important to note that in addition to the metabolic effects, adenosine administration also causes other acute physiological responses including hypothermia and torpor, vasodilation via A2A receptors on vascular smooth muscle, and bradycardia via A1 receptors in the sinoatrial node [57,58]. Reitman and colleagues have shown that peripheral agonism of A2A and A3 and central agonism of A1 and A2B are all sufficient to cause hypothermia in mice. Moreover, this group also showed that agonism of A1, A2A, and A2B were all sufficient to decrease total energy expenditure [59,60]. These changes in activity, temperature, blood pressure, and heart rate could trigger stress pathways and thereby modulate the magnitude and character of the glucose and NEFA responses to adenosine, an added layer of complexity that future work will need to disentangle.

In summary, this study shows that administration of a single bolus of adenosine is sufficient to drive an acute and profound increase in blood glucose and NEFA, with this response enhanced in the postprandial state. Mechanistically, adenosine acts via two adenosine receptors, A2A and A2B in the central nervous system. Our data are consistent with a model whereby activation of these receptors stimulates a sympathetic release of catecholamines into the periphery which then act on α and β adrenergic receptors in the liver and adipose tissue to drive the release of glucose and NEFA. A2A and A2B receptors are individually sufficient, but collectively necessary for mediating this process. That is, stimulation of either receptor is sufficient to promote glucose and NEFA release, but both receptors must be simultaneously inhibited in order to block this effect. We posit that A2A and A2B signaling is highly relevant in modulating postprandial metabolism, and that targeting these signaling pathways may serve as a therapeutic target for mitigating the morbidity of metabolic diseases such as obesity and type II diabetes.

## AUTHOR CONTRIBUTIONS

**DSL**- Writing- original draft, review and editing, investigation, supervision, validation, visualization, data curation, formal analysis. **BDS**- Conceptualization, investigation, methodology, writing- review & editing. **SRH**- Investigation, data curation, conceptualization **JT, RMR, AJN**- Investigation. **MCL, MEG**- Conceptualization. **GKN**-Investigation, methodology. **MPB-** supervision. **SBGA-** Investigation, methodology, supervision, validation. **JL-** Conceptualization, methodology, supervision, writing- review and editing. **TEH-**Conceptualization, formal analysis, funding acquisition, methodology, project administration, supervision, writing- original draft, review and editing.

## ACKNOWLEDGEMENTS

The authors would like to thank Adovate, LLC for their donation of the drugs ATL-844, ADO-313, and BW-1433. All statistical analysis and graphical figures were generated with GraphPad Prism. BioRender was used to generate Figure 8.

**Figure 8:**
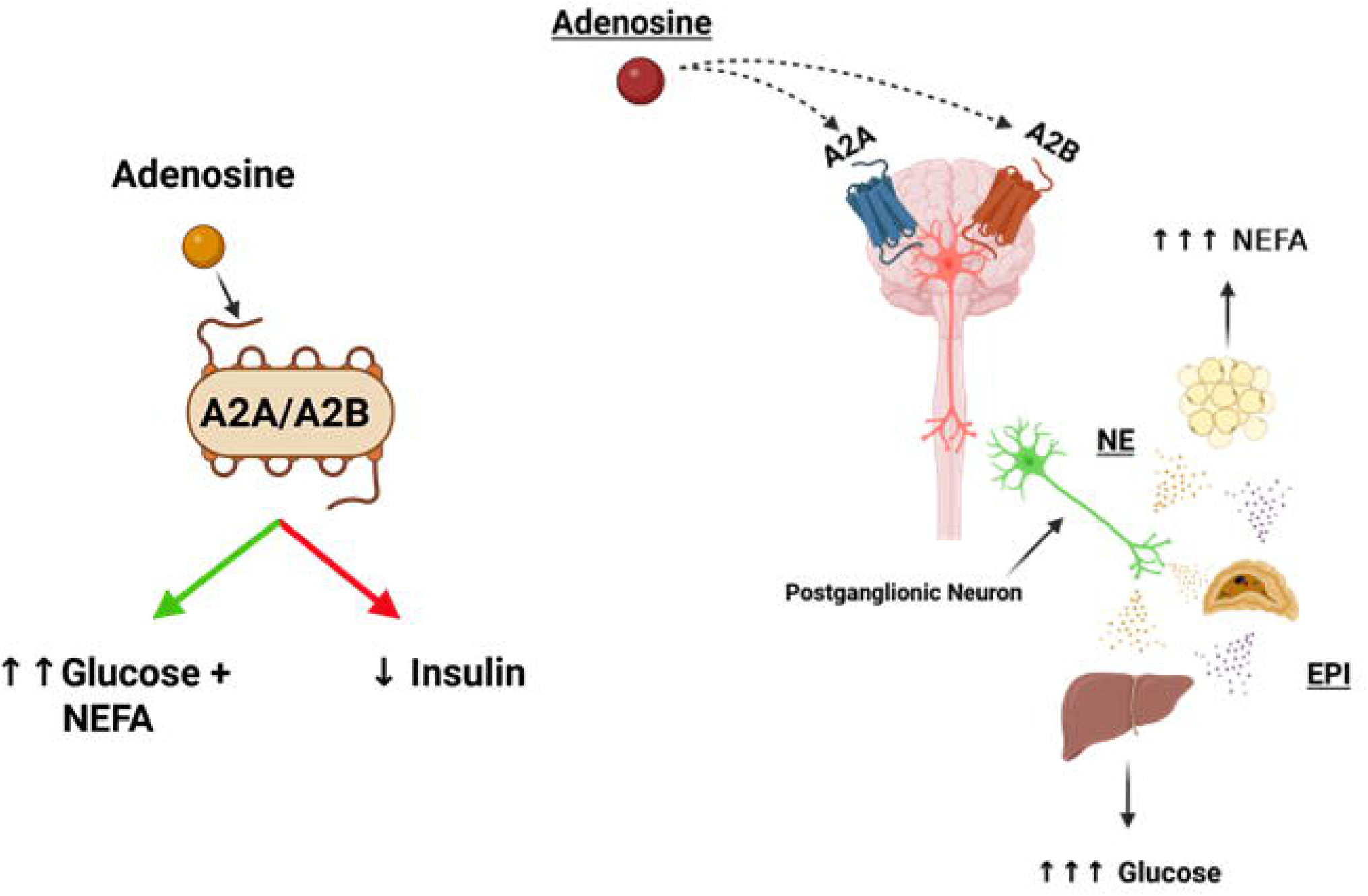
Proposed mechanism of A2A and A2B-stimulated glucose and NEFA excursion: Injected adenosine travels to the brain and activates A2A and A2B receptors in central neurons. This causes an increase in sympathetic signaling into the periphery. Here, postganglionic neurons that innervate into the liver and adipose tissue release norepinephrine (NE) directly onto these tissues to promote hepatic glucose production and lipolysis. Additionally, postganglionic neurons that innervate into the adrenal cortex stimulate a release of epinephrine (EPI), which circulates to the liver and adipose tissue to also promote hepatic glucose production and adipose lipolysis. This effect is enhanced under postprandial conditions due to adenosine-mediated insulin suppression.

