## Supplemental Figures for "Adenosine A2A and A2B receptor signaling in neurons promotes glucose and fatty acid release in the postprandial state"

**Supplementary 1: Adenosine promotes a postprandial glucose and NEFA release in female mice.** WT female mice were fasted and refed as previously described. **(A**) Blood glucose levels of fasted mice i.p. injected with vehicle or adenosine 100 mg/kg. **(C)** Comparison of AUC of blood glucose excursions from A-B. Serum NEFA was measured from the same **(D)** fasted and (**E)** refed female mice. **(F)** Comparison of AUC of the serum NEFA excursions from D-E. Female mice were fasted overnight and then refed as previously mentioned. At t = -30, mice were i.p. injected with either vehicle or a combination of ATL-844 and SCH-58261 (ATL + SCH) (3 mg/kg each). At t = 0, all mice received an additional i.p. injection of adenosine (100 mg/kg). **(G)** Blood glucose and **(H)** Serum NFEA was recorded over 2-hours and AUC of each group was plotted. N"s for each group range from 4-6. Error bars represent SD. *$p < 0.05$, **$p < 0.01$, ***$p < 0.001$, ****$p < 0.0001$. Statistical analyses were done by Welch's t-test (G-H) or two-way ANOVA (C, F).

**Supplementary 2: Individual A2A or A2B inhibition is insufficient to abolish AMP-induced glucose and NEFA excursion.** Refed WT mice were i.p. injected with either vehicle or the A2A inhibitor ZM-241385 (3 mg/kg) at t = -30 minutes. At t = 0 minutes, all mice received an additional i.p. injection of AMP (150 mg/kg). **(A)** Blood glucose and **(B)** serum NEFA were tracked for two hours. Refed WT mice were i.p. injected with either vehicle or the A2B inhibitor ATL-844 (3 mg/kg) at t = -30 minutes. At t = 0 minutes, all mice received an additional i.p. injection of AMP (150 mg/kg). **(C)** Blood glucose and **(D)** serum NEFA were tracked for two hours. N's for each group range from 3-7. Error bars represent SD. *$p < 0.05$, **$p < 0.01$, ***$p < 0.001$, ****$p < 0.0001$. All statistical analyses were done by Welch's T-test.

**Supplemental Figure 3. Confirming double knockout of A2A and A2B in genomic DNA in UBC-A2A+B mice.** Genomic DNA was extracted from the brain tissue from two UBC-A2A$^{fl/fl}$ +A2B $^{fl/fl}$ Cre - (WT) and two UBC-A2A$^{fl/fl}$ +A2B $^{fl/fl}$ Cre + (KO) mice two weeks after tamoxifen injection. **(A)** To show evidence of A2A excision, primers were designed upstream of the 5' loxP site (A2A F) and downstream of the 3' loxP site (A2A R). WT gDNA produces a large 2.2 kB band, and KO gDNA (recombined) produces a smaller <1 kB band. **(B)** Using the same gDNA, this process was repeated for A2B. Primers were designed regions upstream of the 5' loxP site (A2B F) and downstream of the 3' loxP site (A2B R). Due to the large insert in the 3' UTR of A2B$^{fl/fl}$ allele, the WT product is too large to be produced under normal PCR conditions while the KO band (excised) is small enough to be produced to generate a band of 800 bp.

**Supplemental Figure 4: A2A and A2B signaling in adipose tissue and liver do not affect adenosine effects on NEFA and glucose**

A2A$^{fl/fl}$ +A2B $^{fl/fl}$ mice were crossed with the adipoQ-Cre-ERT2 promoter to generate tamoxifen-inducible adipose-specific A2A+A2B double KO mice (FA2A+B). Refed FA2A+B WT and KO mice were given an i.p. bolus of BAY-60-6583 (0.2 mg/kg) and their **(A)** blood glucose and **(B)** serum NEFA were tracked over a five-hour period. A2A$^{fl/fl}$ +A2B $^{fl/fl}$ Cre- mice were given an i.v. injection of AAV8-TBG-Cre or AAV8-TBG-eGFP or to generate liver-specific A2A+A2A DKO mice (AAV-Cre) or WT controls (AAV-eGFP). Refed mice were given a bolus of BAY-60-6583 (0.2 mg/kg) and their **(C)** blood glucose and **(D)** serum NEFA were tracked over a five-hour period. N's for each group range from 5-7. Error bars represent SD. *p < 0.05, **p < 0.01, ***p < 0.001, ****p < 0.0001. All statistical analyses were done by Welch's T-test.

**Supplemental Figure 5: A2A+A2B DKO in peripheral neurons shows no change metabolic change in response to adenosine receptor agonism.** UBC-A2A$^{fl/fl}$ +A2B$^{fl/fl}$ Cre negative mice were tail vein injected with AAV-PHP.S-hSyn1-eGFP-Cre or AAV-hSyn1-tdTomato and given a three-week washout period. Refed AAV-tdTomato and AAV-Cre mice were given an i.p. bolus of adenosine (100 mg/kg) and **(A)** blood glucose and **(B)** serum NEFA were recorded over a two-hour period. Refed AAV-tdToamto and AAV-Cre mice were challenged with an i.p. bolus of the A2B agonist BAY 60-6583 (0.2 mg/kg) and **(C)** blood glucose and **(D)** serum NEFA were recorded over a five-hour period. Refed AAV-tdTomato and AAV-Cre mice were given an i.p. bolus of the A2A specific agonist ATL-313 (0.2 mg/kg) and **(E)** blood glucose and **(F)** serum NEFA were recorded over a three-hour period. **(G)** Superior cervical ganglion were isolated from AAV-Cre-GFP, stained with DAPI, and imaged on a fluorescent microscope to confirm GFP-positive cells. Refed WT mice were given an i.p. injection of either vehicle or BAY (0.02 mg/kg). **(H)** Blood glucose an **(I)** serum NEFA was measured over a two-hour period. N's for each group range from 5 – 9. Error bars represent SD. *$p < 0.05$, **$p < 0.01$, ***$p < 0.001$, ****$p < 0.0001$. All statistical analyses were done by Welch's T-test.

**Supplemental Figure 6**: **Confirmation of Baf53b-A2A+A2B and ROSA26-A2A+A2B double knockout mice.** Genomic DNA was extracted from the brain tissue and liver tissue of one Baf53b-A2A$^{fl/fl}$ + A2B $^{fl/fl}$ WT (Cre -) and one Baf53b-A2A$^{fl/fl}$ +A2B $^{fl/fl}$ DKO (Cre +) mouse. **(A)** PCR was run from brain gDNA (B) and liver gDNA (L) from Baf53b-A2A+B WT and DKO mice using the A2A knockout primers and PCR conditions from Supplemental Figure 3. **(B)** Using the same gDNA, this process was repeated for A2B using the A2B knockout primers from

Supplemental Figure 3. Genomic DNA was extracted liver and brain tissue from one ROSA-$A2A^{fl/fl}$ + A2B $^{fl/fl}$ Cre - (WT) and one ROSA-$A2A^{fl/fl}$ +A2B $^{fl/fl}$ Cre + (DKO) mouse. **(C)** PCR was run from brain gDNA (B) and liver gDNA (L) from ROSA-A2A+B WT and DKO mice using the A2A knockout primers and PCR conditions from Supplemental Figure 3. **(D)** Using the same gDNA, this process was repeated for A2B using the A2B knockout primers and PCR conditions from Supplemental Figure 3.

**Supplemental Figure 7: Adenosine dose not suppress insulin levels in Baf-A2A+B DKO mice.**

Blood insulin levels in refed **(A)** Baf-A2A+B WT and **(B)** Baf-A2A+B DKO mice before (t = 0 min) and after (t = 60 min) an adenosine bolus (100 mg/kg).

### Supplemental Figure 1

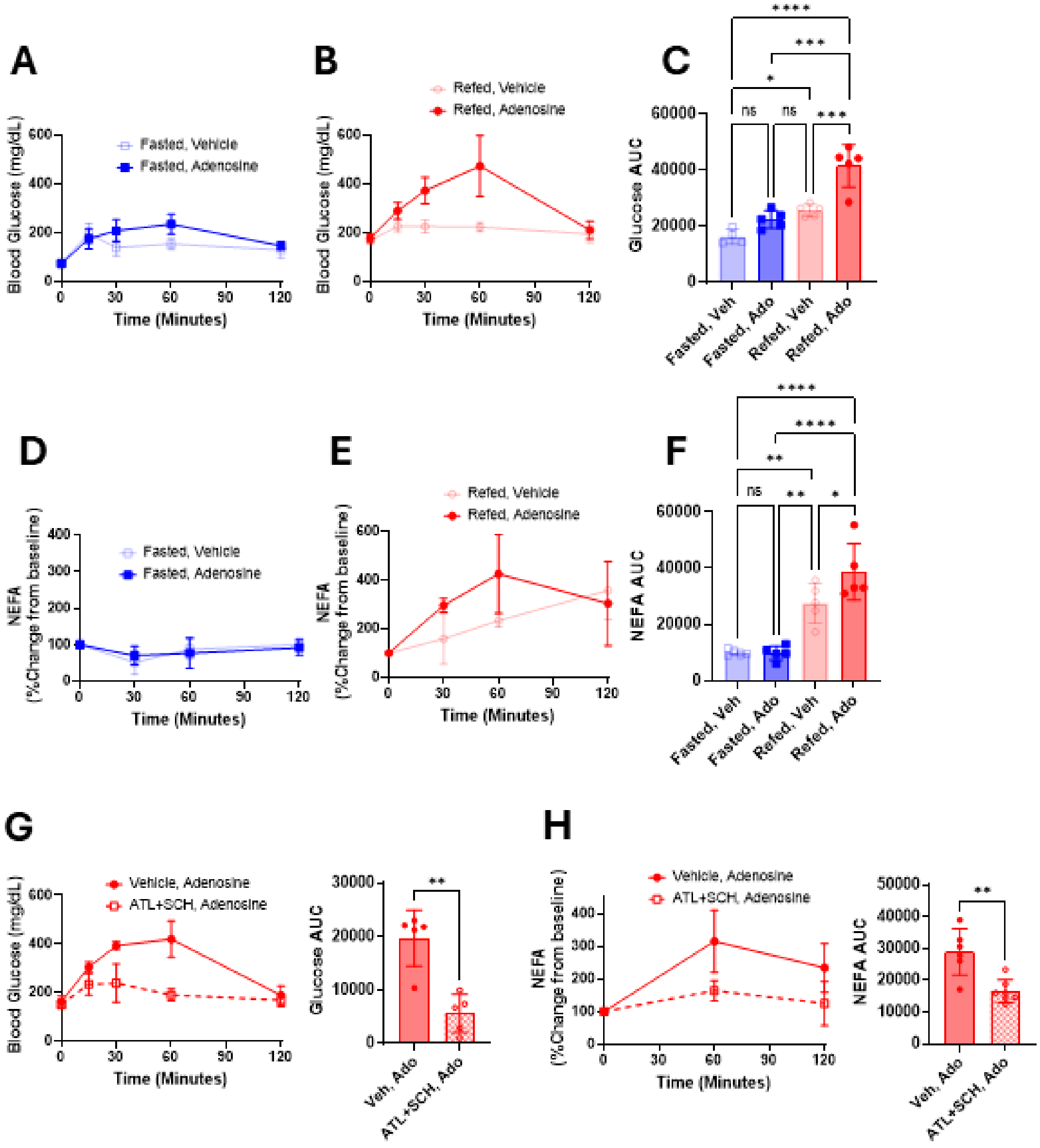

### Supplemental Figure 2

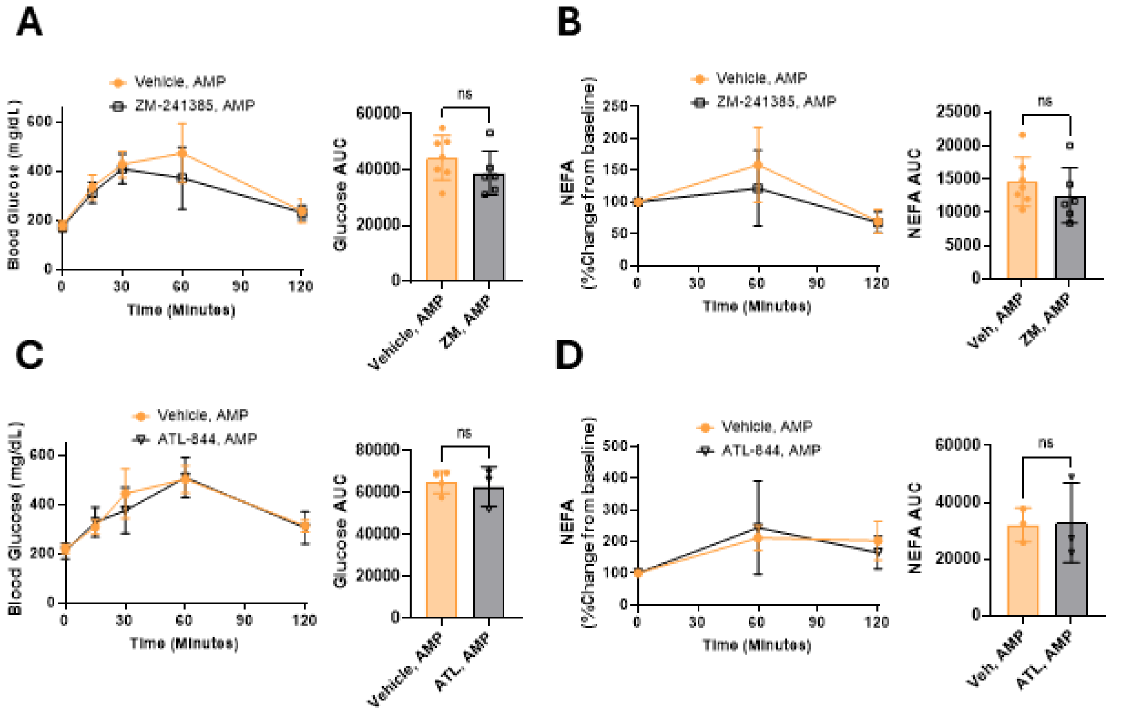

### Supplemental Figure 3

**A**

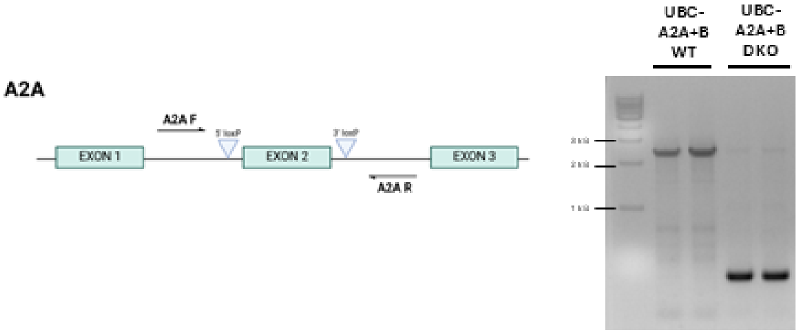


**B**

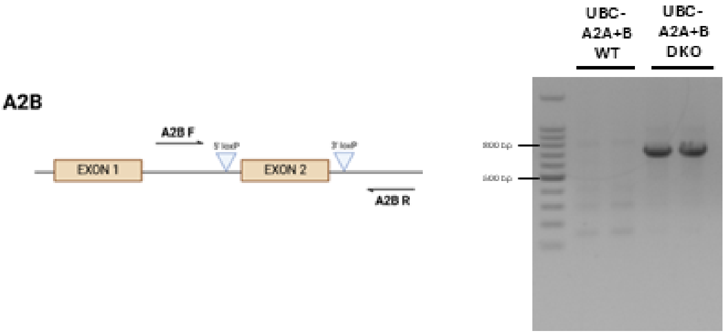

### Supplemental Figure 4

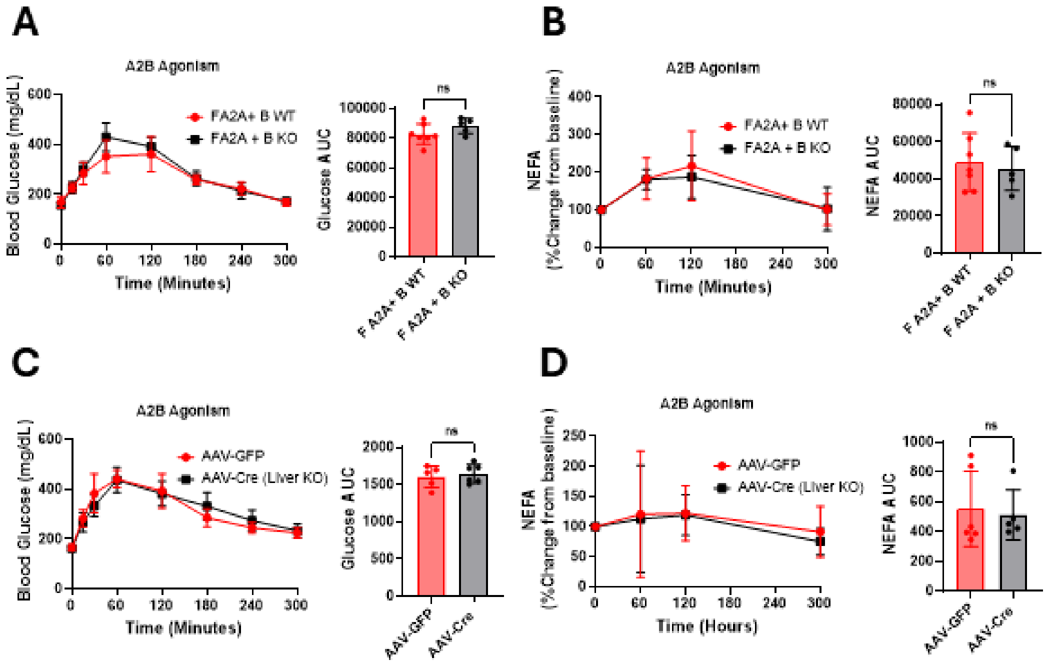

### Supplemental Figure 5

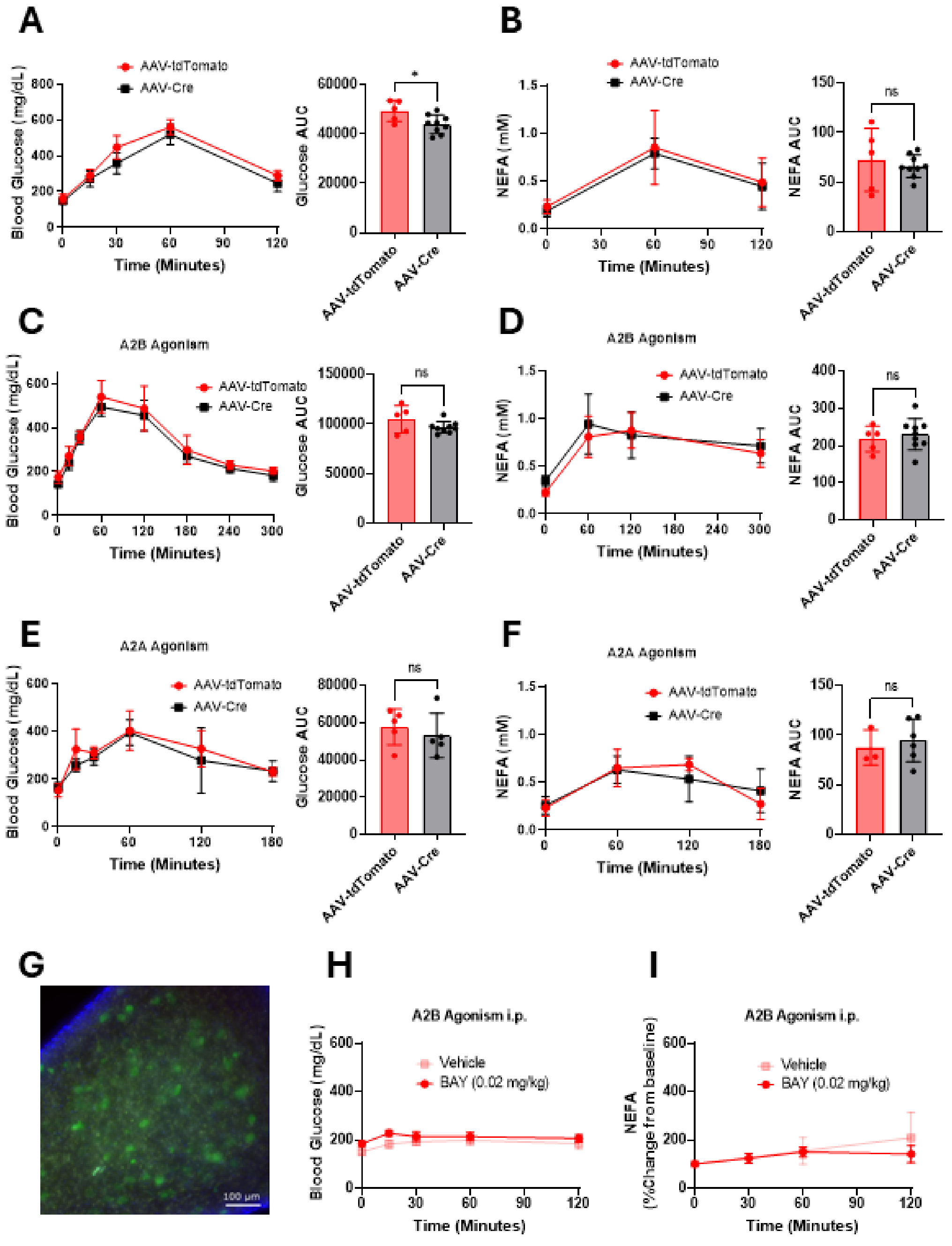

### Supplemental Figure 6

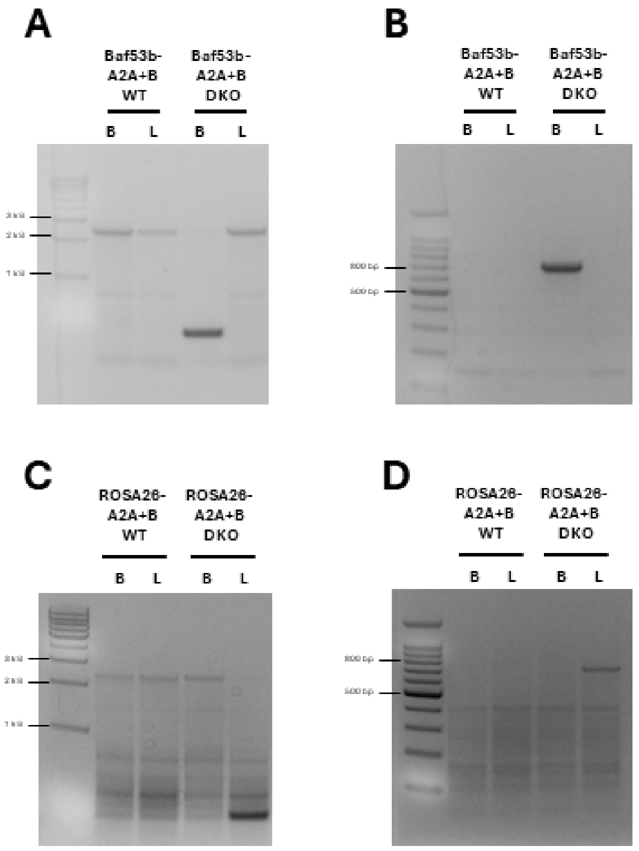

### Supplemental Figure 7

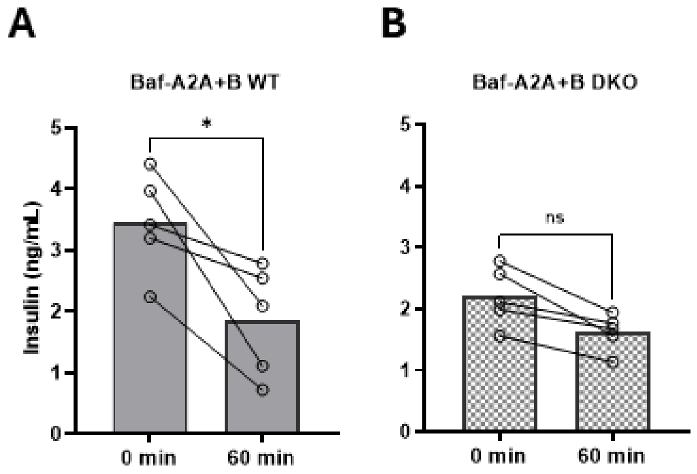
